# A systematic interactome of SET1C expands its functional landscape and identifies candidate regulatory connections

**DOI:** 10.1101/2025.11.23.690026

**Authors:** Pierre Luciano, Kihyun Park, Stéphane Audebert, Luc Camoin, Carlos A. Niño, Da Kyeong Park, Isabella E. Maudlin, Marion Dubarry, Lara Lee, Marlene Oeffinger, Jean D. Beggs, Young Hye Kim, Jaehoon Kim, Bernhard Dichtl, Vincent Géli

## Abstract

Set1 is the catalytic subunit of SET1C or COMPASS, which methylates histone H3K4 and serves as a scaffold for the association of seven tightly bound polypeptides. We have employed yeast two-hybrid screenings to determine the interactome of Set1 and each subunit, providing a unique resource for exploring known and novel roles of the complex. Our screenings identified a multitude of potential interactors involved in chromatin regulation, DNA replication, meiotic breaks, and Ty transposition, processes previously associated with SET1C. Consistent with Set1 being an RNA-binding protein, the screens link SET1C to multiple aspects of RNA biogenesis, including pre-mRNA splicing and polyadenylation. The results reveal that several importins are candidate interactors of Set1, along with RGG motif-containing proteins, providing insights into the mechanisms by which Set1 moves between cytoplasmic and nuclear compartments. We further reveal that reconstituted SET1C interacts with the AT hook domain of the chromatin remodeler Snf2 and methylates multiple arginines within this domain. *In vivo*, we report that the ARTSTRGR AT-hook motif is methylated in a Set1-dependent manner revealing new interplay between lysine and arginine methylation.

## INTRODUCTION

Post-translational modifications of histones shape the chromatin landscape and provide essential mechanisms of regulating DNA accessibility, thereby controlling gene expression, genome maintenance and transmission. Multiprotein complexes of the SET/MLL family methylate histone H3 on lysine 4 (H3K4), which modulates multiple aspects of genome biology (Ruthenburg *et al*, 2007). Each complex contains additional factors specifying their recruitment to explicit chromatin domains and their specific biological effects (Cenik & Shilatifard, 2021). The budding yeast Set1 complex called COMPASS (for Complex of Proteins Associated with Set1) or SET1C has proved to be an excellent model to study the SET1/MLL family complexes. In *Saccharomyces cerevisiae*, all H3K4 methylation is carried out by SET1C that is composed of Set1, the catalytic subunit, acting as a scaffold for seven other components (Swd1 [RbBP5], Swd2 [WDR82], Swd3 [WDR5], Bre2 [ASHL2], Sdc1 [DPY30], Spp1 [CFP1] and Shg1 [BOD1]) (Nagy *et al*, 2002; Briggs *et al*, 2001; Roguev *et al*, 2001; Miller *et al*, 2001). Swd1, Swd3, Bre2 and Sdc1 copurify with the SET domain of Set1, yet none of these proteins can interact alone with the SET domain, suggesting that interactions between subunits are required for catalytic domain formation (Kim *et al*, 2013). The crystal structure of the SET domain of Set1 associated with Swd1, Swd3, Bre2, Sdc1 shows that the catalytic module is organized by Swd1, whose C-terminal tail nucleates Swd3 and a sub-complex formed by Bre2-Sdc1 (Qu *et al*, 2018; Hsu *et al*, 2018). For its side, Spp1 associates with the N-SET domain of Set1, while Shg1 binds to the central region of Set1 and Swd2 contacts the N-terminal region of Set1 (Kim *et al*, 2013; Roguev *et al*, 2001; Dehe *et al*, 2006; Halbach *et al*, 2009). Swd1 contacts Spp1 and also interacts with the N-terminal region of Set1, suggesting that the C- and N-terminal regions of Set1 may interact with each other (Qu *et al*, 2018; Acquaviva *et al*, 2013b; Jeon *et al*, 2018). Such SET1C organization was confirmed by cryo-electron microscopy and cross-linking experiments performed on the full-length complex (Wang *et al*, 2018). Interestingly, SET1C has a remarkable mode of assembly that is initiated in the cytoplasm while the nascent Set1 polypeptide emerges from the ribosome. Set1 is initially bound during its translation, by Shg1, Spp1 and Swd1, then Swd2, Swd3, Bre2 and Sdc1 associate with the initial pre-complex to form the full SET1C. This explains why Set1 is associated with its own mRNA (Luciano *et al*, 2017; Halbach *et al*, 2009).

Swd2, the only essential SET1C subunit, is also part of the APT complex (for Pta1-associated), a subcomplex of the Cleavage Polyadenylation Factor (CPF) involved in the transcriptional termination of mRNA and snoRNA suggesting a functional link between SET1C and 3’ end formation/termination (Cheng *et al*, 2004; Nedea *et al*, 2003; Dichtl, 2004). Set1 has also been linked genetically to the Nrd1-Nab3-Sen1 (NNS) complex (Kim *et al*, 2006; Arigo *et al*, 2006), but intriguingly, it was reported to have two seemingly contradictory effects on ncRNA termination: on the one hand Set1 was suggested to promote efficient snoRNA termination mediated by Nrd1 (Terzi *et al*, 2011), and on the other hand it appeared to interfere with the early termination of a large number of ncRNAs (Castelnuovo *et al*, 2014; Margaritis *et al*, 2012). Interestingly, various lysines distributed between Nrd1, Nab3 and Sen1, are methylated, notably lysine 363 in the RNA Recognition Motif (RRM) of Nab3 whose mono methylation depends on Set1 and Set3 (Lee *et al*, 2020).

Set1 interacts co-transcriptionally with the RNA polymerase II Carboxy Terminal Domain (PolII CTD) phosphorylated on Ser5 of the heptad repeats, producing an H3K4 methylation gradient that starts at nucleosome +1 and fades away from the promoter (Soares *et al*, 2017; Ng *et al*, 2003). It was recently shown that the N-terminal region of Set1 and Swd2 interact cooperatively with the CTD (Carboxy Terminal Domain) of Rbp1 (RNA polymerase II large subunit) promoting SET1C recruitment to transcription elongation complexes at the 5′ ends of genes (Bae *et al*, 2020). Deletion of residues 200-210 of Set1 abolishes interaction with the Rpb1-CTD and Swd2, while deletion of the first 200 amino acids of Set1 strongly reduces trimethylation of H3K4 (H3K4me3) at the 5’ end of transcribed genes (Bae *et al*, 2020). Interestingly, Swd2 is ubiquitinated by Rad6/Bre1 and preventing Swd2 ubiquitination affects H3K4 trimethylation with a concomitant reduction in Spp1 recruitment to chromatin suggesting a cross-talk between Swd2 and Spp1(Vitaliano-Prunier *et al*, 2008). In a similar vein, di- and tri-methylation of H3K4 by Set1, which depends on prior ubiquitination of histone H2B by Rad6/Bre1 (Dover *et al*, 2002; Sun & Allis, 2002), requires a contact between Spp1 and Swd1 (Hsu *et al*, 2019; Jeon *et al*, 2018). Other modes of SET1C recruitment to chromatin must exist in the absence of the Set1 N-terminal region, probably via direct interactions between SET1C’s catalytic domain and the nucleosome (Jeon *et al*, 2018; Dehé & Géli, 2006; Thornton *et al*, 2014).

The central region of Set1 contains two RRMs positioned in tandem (Trésaugues *et al*, 2006) through which Set1 binds directly to RNA. This binding contributes to retain Set1 in the 5’ region of genes thus facilitating their H3K4 trimethylation (Luciano *et al*, 2017; Battaglia *et al*, 2017; Sayou *et al*, 2017). Both the N-SET domain and Spp1 also contribute to Set1 binding to RNA. Set1 associates post-transcriptionally with transcripts produced by specific classes of genes including snRNA, Ty1 and a number of genes involved in adaptive responses (Luciano *et al*, 2017). The dRRM itself is flanked by an autoinhibitory region that negatively regulates H3K4me3 (Schlichter & Cairns, 2005). It is possible that in the context of full-length Set1, the alternative mode of SET1C recruitment may be inhibited by the autoinhibitory domain.

In *S. cerevisiae*, H3K4me3 methylation is counteracted by the demethylase Jhd2, a conserved JARID1 family protein (Huang *et al*, 2010; Liang *et al*, 2007). The erasure of ancestral histone methylation states results from both active enzymatic demethylation by Jhd2 and passive dilution of parental histones during replication (Radman-Livaja *et al*, 2010). SET1C is a dimer and its dimerization depends on the Sdc1 subunit (Choudhury *et al*, 2019). It has been proposed that the symmetrical methylation of H3K4 on nucleosomes is a consequence of the dimeric nature of SET1C and that Jhd2 preferentially demethylates asymmetrical H3K4me3 (Choudhury *et al*, 2019). Interestingly, H3K4me3 at environmental stress genes depends on H3-P16 isomerisation, a process that controls K4me3 by balancing the actions of Jhd2 and Spp1 (Howe *et al*, 2014).

The complexity of the *S. cerevisiae* SET1C, the specific arrangement of subunits within the complex, and the specific roles of certain subunits, notably Spp1 and Swd2 (Acquaviva *et al*, 2013a), raise the question of the contribution of SET1C and of its subunits to multiple processes. Overall, H3K4 methylation, SET1C and its individual subunits have been involved in multiple processes in *S. cerevisiae* such as meiotic recombination (He *et al*, 2019; Sollier *et al*, 2004; Borde *et al*, 2008; Acquaviva *et al*, 2013b; Sommermeyer *et al*, 2013; Karányi *et al*, 2018; Adam *et al*, 2018), stress response and epigenetic transcriptional memory (D’Urso *et al*, 2016; Kim & Buratowski, 2009; Weiner *et al*, 2012; Deshpande *et al*, 2022; Nadal-Ribelles *et al*, 2015)), DNA repair (Faucher & Wellinger, 2010), telomere and rDNA silencing (Jezek *et al*, 2023; Corda *et al*, 1999; Briggs *et al*, 2001; Bryk *et al*, 2002; Santos-Rosa *et al*, 2004; Nislow *et al*, 1997), cell wall biogenesis (Nislow *et al*, 1997), chromosome segregation (Beilharz *et al*, 2017; Zhang *et al*, 2000), antisense transcription (Murray *et al*, 2015; van Dijk *et al*, 2011; Margaritis *et al*, 2012; Castelnuovo *et al*, 2014), transcription termination (Kaczmarek Michaels *et al*, 2020; Nedea *et al*, 2003; Dichtl, 2004; Cheng *et al*, 2004; Nedea *et al*, 2008; Terzi *et al*, 2011; Soares & Buratowski, 2012; Castelnuovo *et al*, 2013; Lee & Wang, 2018), Ty silencing (Luciano *et al*, 2017; Berretta *et al*, 2008), chronological aging (Gong *et al*, 2023; Walter *et al*, 2014; Mei *et al*, 2019), ergosterol homeostasis (South *et al*, 2013), lipid homeostasis (Giaever *et al*, 2019), and DNA replication (Ghaddar *et al*, 2023; Sollier *et al*, 2004; Rizzardi *et al*, 2012; Chong *et al*, 2020; Delamarre *et al*, 2020; Santos-Rosa *et al*, 2021; de La Roche Saint-André & Géli, 2021; Serra-Cardona *et al*, 2022).

The multiple roles of Set1 and its subunits led us to perform global two-hybrid screening to identify interactors either of Set1-full length, or of its N- and C-terminal regions and of each of the individual subunits (Swd2, Shg1, Spp1, Swd1, Swd2, Sdc1, Bre2). The identification of interactors is discussed not only for each bait, but also as a whole, revealing new potential functions for the complex. In addition, we demonstrate that the Snf2 AT-hook is methylated by SET1C *in vitro*. In vivo, deleting *SET1* abrogates arginine methylation within the ARTSTRGR motif of Snf2 AT-hook. These results reveal new interplay between Lysine and arginine methylation. This work is an invaluable resource for further exploring known and unsuspected roles of the SET1C complex and its subunits.

## RESULTS

### Identification of Set1 and Set1 subunits interactors

We produced a total of ten yeast two-hybrid (Y2H) screens (Hybrigenics, See Methods). For the Set1 protein, we performed a total of three screens using as bait the full-length Set1 protein (Set1 FL), the amino-terminal region including amino acids 1-754 or a C-terminal fragment including amino acids 754-1081 (Fig. 1A). We have chosen to separate Set1 into these 2 regions because they have been described as having well-defined properties. The Set1 1-754 fragment includes the domain involved in Set1 recruitment to chromatin, the double RRM (Trésaugues *et al*, 2006), and the central self-inhibitory domain (Schlichter & Cairns, 2005) that all have regulatory roles (Dehé & Géli, 2006). The Set1 region 754-1081 contains the N-SET domain, the SET domain and the post-SET domain: the latter is capable of methylating H3K4 on its own (Thornton *et al*, 2014) (Fig. S1). For each of the subunits (SU) of the complex whose organization is described in Fig. 1A we used the whole protein as bait. Each gene encoding Set1 (and its fragments) or its subunits was fused to the C-terminus the Gal4-BD except for Swd2 that was fused to the N-terminus of Gal4-BD. These screens have proven their power and effectiveness. In particular, they previously identified Mer2 as an interactor of Spp1 (Acquaviva *et al*, 2013b), and the CTD of Rpb1 (Rpo21 in Table S2) as an interactor of the N-terminal region of Set1 (Bae *et al*, 2020). The results of the ten Y2H screens are presented in Table S2. This table also highlights interactors that are shared among multiple subunits as well as those common to different Set1 fragments. Y2H interactions indicate potential protein–protein associations but do not constitute definitive proof of interaction, accordingly the list of candidates may include false positives. Each interactor is characterised with a confidence score based on several criteria, including the frequency with which a prey protein is found for a particular screen and the presence of overlapping fragments, which allow the delineation of the interaction domain involved (see Methods). Very high confidence interactors (indicated by a color code) are likely to interact directly with their bait. For Set1 and its N- and C-terminal fragments, the high confidence Y2H interactors are shown in Fig. 1B. The predicted protein-protein interactions within the Set1 interactors reveal functional information that is detailed in the following paragraphs. The high confident interactors of the seven SET1C subunits are shown in Fig. 1C-E. We found that Spp1, Shg1 and Swd2 interact alone with Set1 (Fig. 1C). The minimum Set1 region for which an interaction is found for each of these 3 subunits is shown in Fig. 1C. The high confidence interactors of the seven SET1C subunits are shown in Fig. 1C-E. We found that Spp1, Shg1 and Swd2 display Y2H interactions with Set1 (Fig. 1C). The high confidence interactors of Spp1, Shg1 and Swd2 are indicated in Fig. 1D (see also Table S2). In contrast, we did not find any interaction between Swd1, Swd3, Bre2 and Sdc1 (SET-c components) and Set1 in the various Y2H screens, suggesting that these subunits must cooperate to bind to Set1. Only Bre2 was found to interact with Sdc1 in both Sdc1 and Bre2 Y2H screens (Fig. 1E). These results are consistent with the structure of the extended SET1C catalytic module and full-length SET1C (Wang *et al*, 2018; Hsu *et al*, 2018; Qu *et al*, 2018).

**Figure 1.**
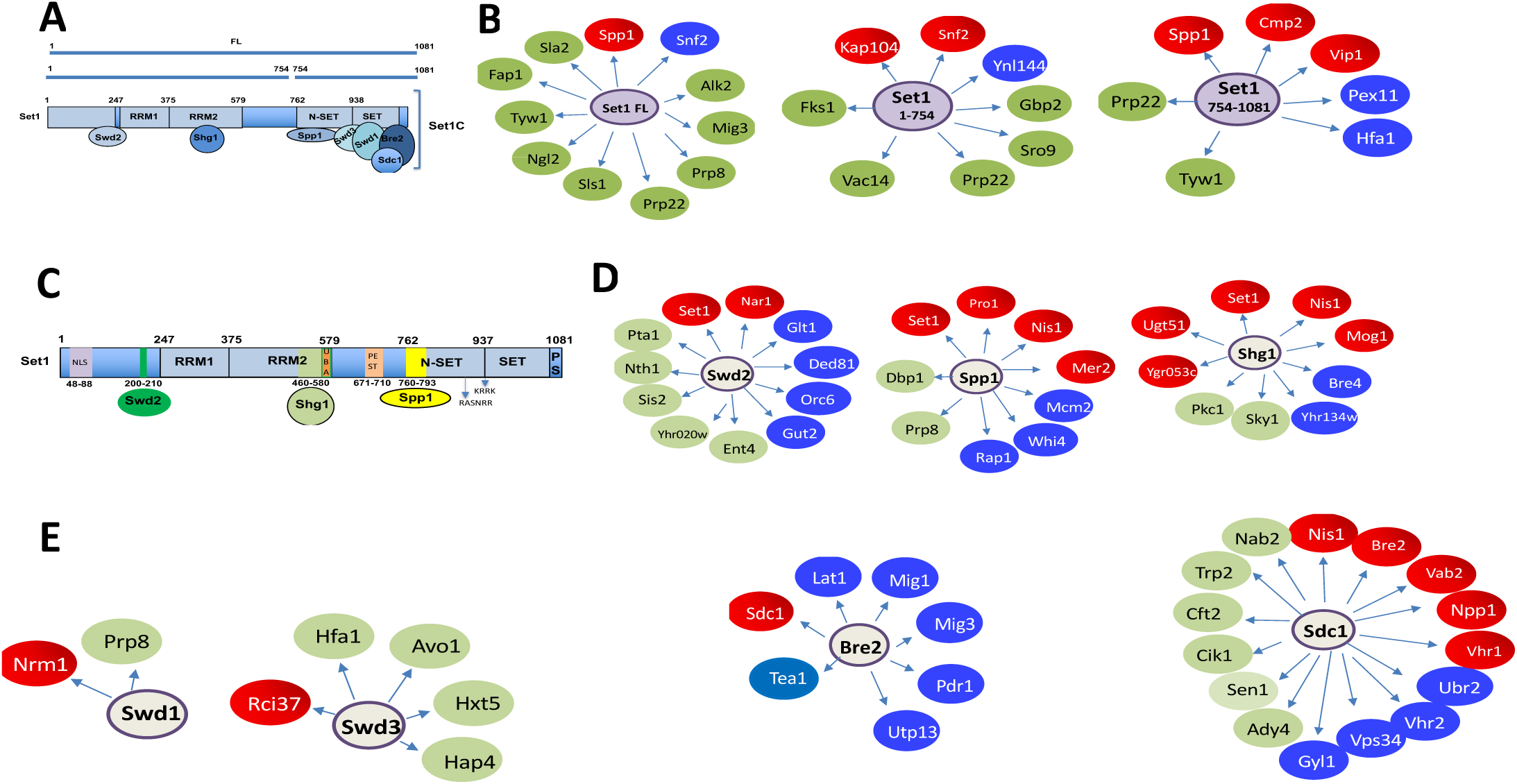
Schematic representation of the Set1 and subunit major interactors identified in the systematic yeast two-hybrid screens. **A**) Schematic representation of Set1 FL and Set1 fragments 1-754 and 754-1081. **B**) Set1 FL, 1-754 and 754-1081 major Y2H interactors). **C**) Interacting regions of Set1 with Swd2, Shg1 and Spp1. **D**) Swd2, Spp1, Shg1 major Y2H interactors. **E**) Swd1, Swd3, Bre2, Sdc1 major Y2H interactors. The term “interactor” is used to mean a high confidence two-hybrid interaction, with the limitations that this entails. The color reflects the Predicted Biological Score (see METHODS). Red, highest confidence; Blue, high confidence; Green, good confidence.

### Set1 1-754 interacts with the importin Kap104

We found that Set1 1-754 interacted with very high confidence with the importin Kap104, suggesting a direct interaction (Fig. 2A). Kap104 has been reported to recognize specific cargos containing a specific class of NLS termed PY-NLS (Soniat *et al*, 2013; Xu *et al*, 2010). This PY-NLS includes a N-terminal or central hydrophobic or basic motif, which often contains hydrophobic (R/H/K) residues and which can include RGG repeats (arginine-glycine-glycine motifs) and a C-terminal Proline-Tyrosine (PY) dipeptide near the C-terminus. Such cargos include the mRNA export factor Nab2, the subunit of the THO/TREX complex Hrp1, and the transcription factor Tfg2 (Lee & Aitchison, 1999; Süel *et al*, 2008). Interestingly, Nab2 is a confident interactor of Sdc1 (Fig. 2A). The Set1 interacting domain (SID) of Kap104 extends from residue 359 to 621 and includes the HEAT-like repeat (Yoshimura & Hirano, 2016) (Fig. 2B). Consistent with this result, we identified two PY-NLS in the N-terminal region of Set1 that are likely recognized by Kap104 (Fig. 2C). To strengthen this observation, we performed AlphaFold modelling of a seven subunit Set1C (Set1-Bre2-Sdc1(x2)-Swd1-Swd3-Spp1) and Kap104 (Fig. 2D). Obtained models showed prediction parameters which are indicative of an overall predicted fold for the complex that was at the threshold for a true structure (pTM = 0.53) and below the confidence threshold for predicting the relative positions of the subunits within the complex (ipTM = 0.5). Analysis of a model in Pymol revealed Kap104 SID binding to Set1-754 as predicted from the Y2H (in the structure we only show the Kap104 SID, while the remaining Kap104 sequences are hidden such that the Set1 PY-NLS is visible). In agreement with Y2H data the model recapitulates Kap104 SID binding to the PY-NLS sequence of Set1 at position 40 - 90; the second PY-NLS of Set1 is not visible in the shown structure and is not interacting with Kap104 in this model. Moreover, the Spp1 and Sdc1-dimer bind on opposing ends of the complex as expected (Qu *et al*, 2018). Taken together the AlphaFold model recapitulates key predictions for the Set1-Kap104 Y2H interaction.

**Figure 2.**
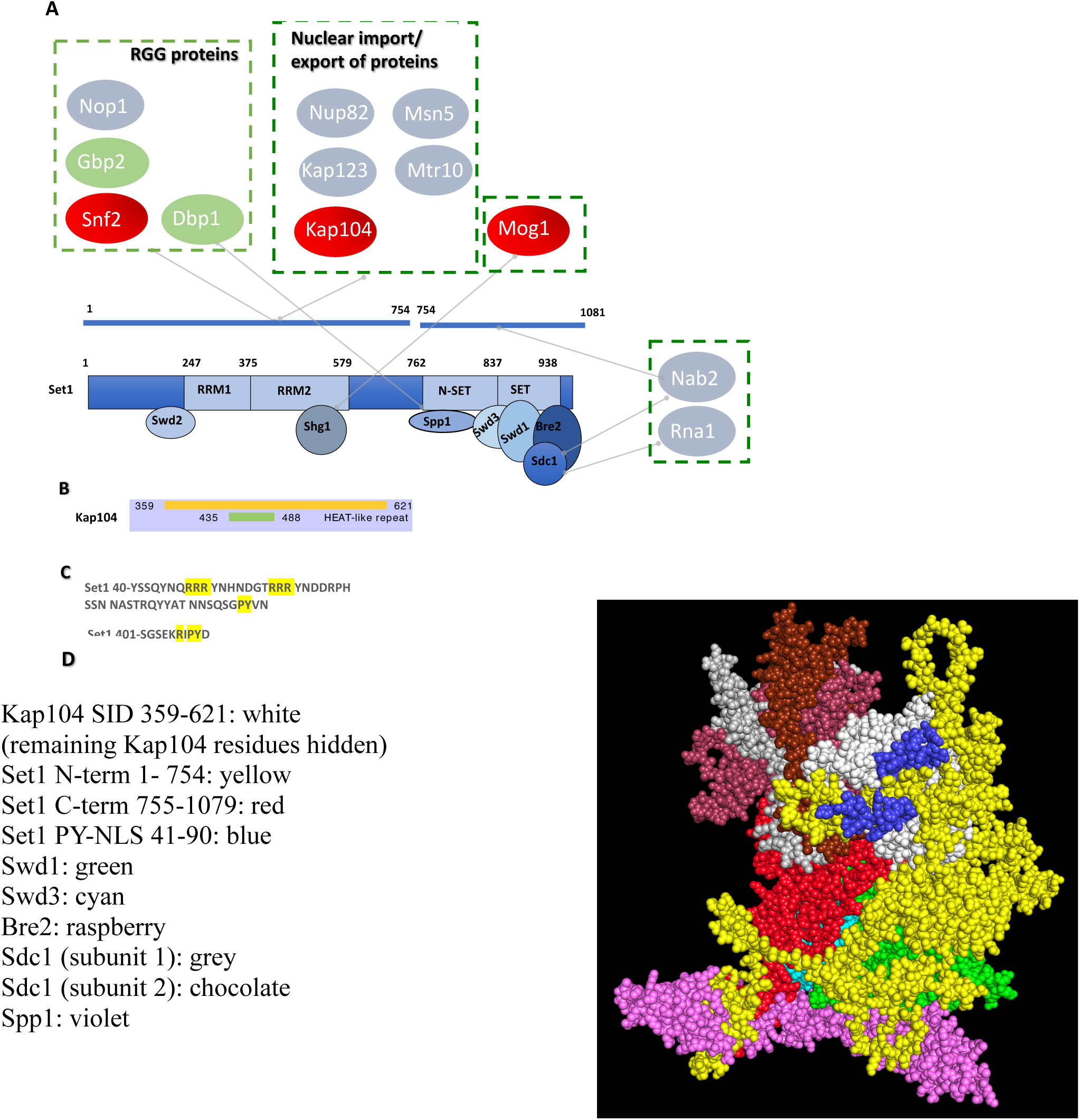
Set1 1-754 interacts with RGG proteins and the importin Kap104. **A**) RGG proteins and import/export proteins interacting with Set1 1-754, Set1 754-1081 and Spp1, Shg1, and Sdc1. **B**) Set1 interacting domain (SID) (blue) within Kap104. Heat like repeat 9 and 10 are represented in purple **C**) PY-NLS in the N-terminal region of Set1. **D**) AlphaFold modelling of a seven subunit Set1C (Set1-Bre2-Sdc1(x2)-Swd1-Swd3-Spp1) and Kap104. A representative model is shown using the following colour code: Kap104 SID 359-621 (white, the remaining Kap104 residues are hidden. Set1 N-term 1-754 (yellow), Set1 C-term 755-1079 (red); Set1 PY-NLS 41-90 (blue); Swd1 (green); Swd3 (cyan); Bre2 (raspberry); Sdc1 (subunit 1, grey); Sdc1 (subunit 2, chocolate); Spp1 (violet).

Shg1, which binds RRM2, interacted with very high confidence with Mog1 (Oliete-Calvo *et al*, 2018). Mog1 has been involved in the modulation of the nucleotide state of Ran-GTP in the nucleus and of Ran-GDP in the cytoplasm, thereby conferring directionality to the nuclear import pathway (Baker *et al*, 2001). Mog1 has been reported to interact directly with the Ran homologue Gsp1 (Oki & Nishimoto, 1998) (Fig. 2A). We have also identified, Kap123, Msn5 (Kap142), Nup82 and Mtr10 as interactors of Set1 1-754 that are also involved in protein import/export (Fig. 2A). Kap123 is a major karyopherin that recognizes NLS of cytoplasmic H3 and H4 (An *et al*, 2017) while Msn5/Kap142 was shown to mediate the import into the nucleus of the subunits of RPA (Yoshida & Blobel, 2001). Nup82 for its part belongs to a module at the cytoplasmic face of the NPC and interacts with karyopherins (Beck & Hurt, 2017) (Fig. 2A). Finally, Mtr10 has been implicated in the nuclear import of the mRNA-binding protein Npl3 (Senger *et al*, 1998; Pemberton *et al*, 1997). Collectively, these multiple interactions reveal insights for understanding the nuclear import of Set1, which was reported to bind co-translationally with Shg1, Swd1, and Spp1 (Halbach *et al*, 2009).

### Set1 1-754 interacts with Snf2 and RGG motif proteins

Set1 1-754 and Set1 FL interacted in the 2H screens with very high confidence with Snf2, the catalytic subunit of the SWI/SNF chromatin remodeling complex (Côté *et al*, 1994) (Fig. 1B, Table S2). Interestingly, two additional Snf2 complex components, Snf5 and Swi1, have been identified in the Set1FL screen (Table S2). Together these observations strongly support the idea that the suggested interaction of Set1 and SWI/SNF is of biological significance. The various Y2H screens revealed Set1 and its subunits interacted with a number of proteins involved in chromatin structure regulation although the relevance of these interactions remains to be demonstrated (Table S2, Fig. S2). Interestingly, the SID region of Snf2 is juxtaposed to an RGG motif composed of several RGG/RG repeats that is often found in RNA-binding proteins (Thandapani *et al*, 2013) and may be the substrate for arginine methyltranferases (McBride *et al*, 2005). RGG repeats are multifunctional motifs that mediate RNA and DNA binding and nuclear import. They have been involved in RNA metabolism and chromatin dynamics possibly via arginine methylation, which modulates RNA affinity and nuclear localization (108). Along the same line, Set1 1-754 also interacted with the mRNA export factor Gbp2 (Poornima *et al*, 2021) and the nucleolar protein Nop1, both of which contain an RGG motif (Fig. S3). Other RGG proteins such as polyadenylated RNA-binding protein Nab2 and the RNA-helicase Dbp1 interacted with Set1 754-1081 (and Sdc1) and Spp1, respectively (Fig. S3). For each of these proteins, the putative SID includes the RGG motif. Of note, both Nop1 and Nab2 are methylated by the Arginine methyltransferase Hmt1 (Smith *et al*, 2020; Green *et al*, 2002).

### SET1C is involved in multiple aspects of RNA biogenesis

Set1 was previously shown to bind RNA nascent transcripts through its dRRM contributing to position Set1 and H3K4me3 predominantly to the 5′ regions of genes. Of note, Set1 showed a higher occupancy within introns, at transcripts from ribosomal DNA (rDNA), and tRNAs (Luciano *et al*, 2017; Battaglia *et al*, 2017; Sayou *et al*, 2017). Moreover, Set1 also binds post-transcriptionally to Ty1 retrotransposon transcripts and mRNA encoding specific transcription factors genes (Luciano *et al*, 2017). The SET1C Y2H interactome revealed that many proteins involved in RNA biogenesis were candidate interactors, interacting directly or indirectly with Set1 or its subunits (Fig. 3). It is not feasible to validate all of these interactions within the limits of this manuscript, and their validity should therefore be interpreted with caution. Nonetheless, these findings provide a useful basis for future research. We find a strong connection between SET1C and pre-mRNA processing, as highlighted by the identification of multiple spliceosome subunits (Fig. 3). Some spliceosome subunits were found in several screens (Fig. 1B, D; Table S2). For instance, the Prp8 subunit of the U5 snRNP that function in critical molecular rearrangements during the splicing process (Grainger & Beggs, 2005) interacted with Set1 FL, Spp1, and Swd1. Similarly, Prp22 interacts with Set1FL and Set1 754-1080. Prp22 is a DEAH-box helicase that associates with newly spliced mRNA and promotes its release from the spliceosome (Will & Lührmann, 2011).

**Figure 3.**
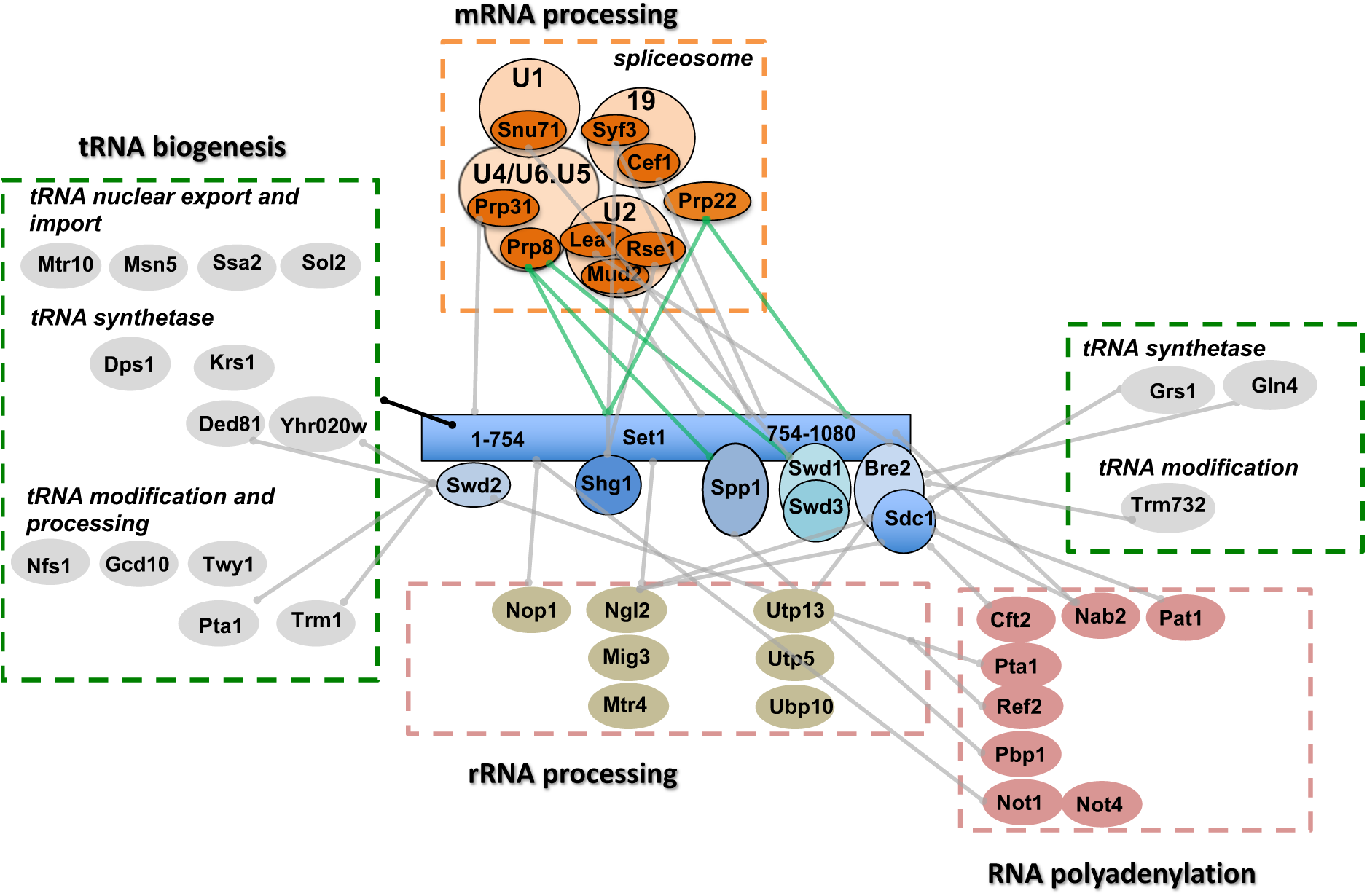
The SET1C Y2H interactome identifies proteins involved in RNA biogenesis. All Y2H interactors are described in Table S2. The processes linked to RNA metabolism in which the different interactors are involved are shown in the figure. The green lines linking Prp22 to Set1FL/Set1 754-1080 and Prp8 to Set1 FL and Spp1 indicate interactions with a high degree of confidence.

We have further strengthened the link between Set1 and Prp22 by showing that Set1 is co-immunoprecipitated with Prp22 *in vivo* in an RNA-independent manner (Fig. 4). The preferential binding of Set1 to genes with introns (Luciano *et al*, 2017) and its interaction with splicing factors, in particular Prp22, suggest that Set1 may be involved in late splicing events. Alternatively, Prp22 and/or other splicing factors could be involved in H3K4 methylation. We also found a number of factors involved in rRNA processing that could be related to the binding of Set1 to ncRNA transcripts derived from the rDNA intergenic spacer regions (Sayou *et al*, 2017) (Fig. 3). Next, and consistent with the observation that SET1C regulates the choice of the polyadenylation site and the recruitment of the cleavage/polyadenylation complex (Kaczmarek Michaels *et al*, 2020), the Y2H screens revealed multiple potential interactions between SET1C and factors involved in RNA polyadenylation (Fig. 3). Some of these interactions are known, such as the interactions between Swd2 with Pta1 and Ref2, all three proteins belonging to the APT complex (Nedea *et al*, 2008). Interestingly, we found that Not1 and Not4 that belongs to the ubiquitin-protein ligase CCR4-NOT (Liu *et al*, 2001) were potential interactors of Set1 1-754 (Fig. 3). This complex was shown to be involved in the regulation of H3K4me3 via a ubiquitin-dependent pathway (Mulder *et al*, 2007; Laribee *et al*, 2007) that was subsequently linked to Jhd2 degradation (Huang *et al*, 2010; Mersman *et al*, 2009). These results raise the question of which of the Y2H interactions described in this study are linked to the regulation of H3K4 methylation states.

**Figure 4.**
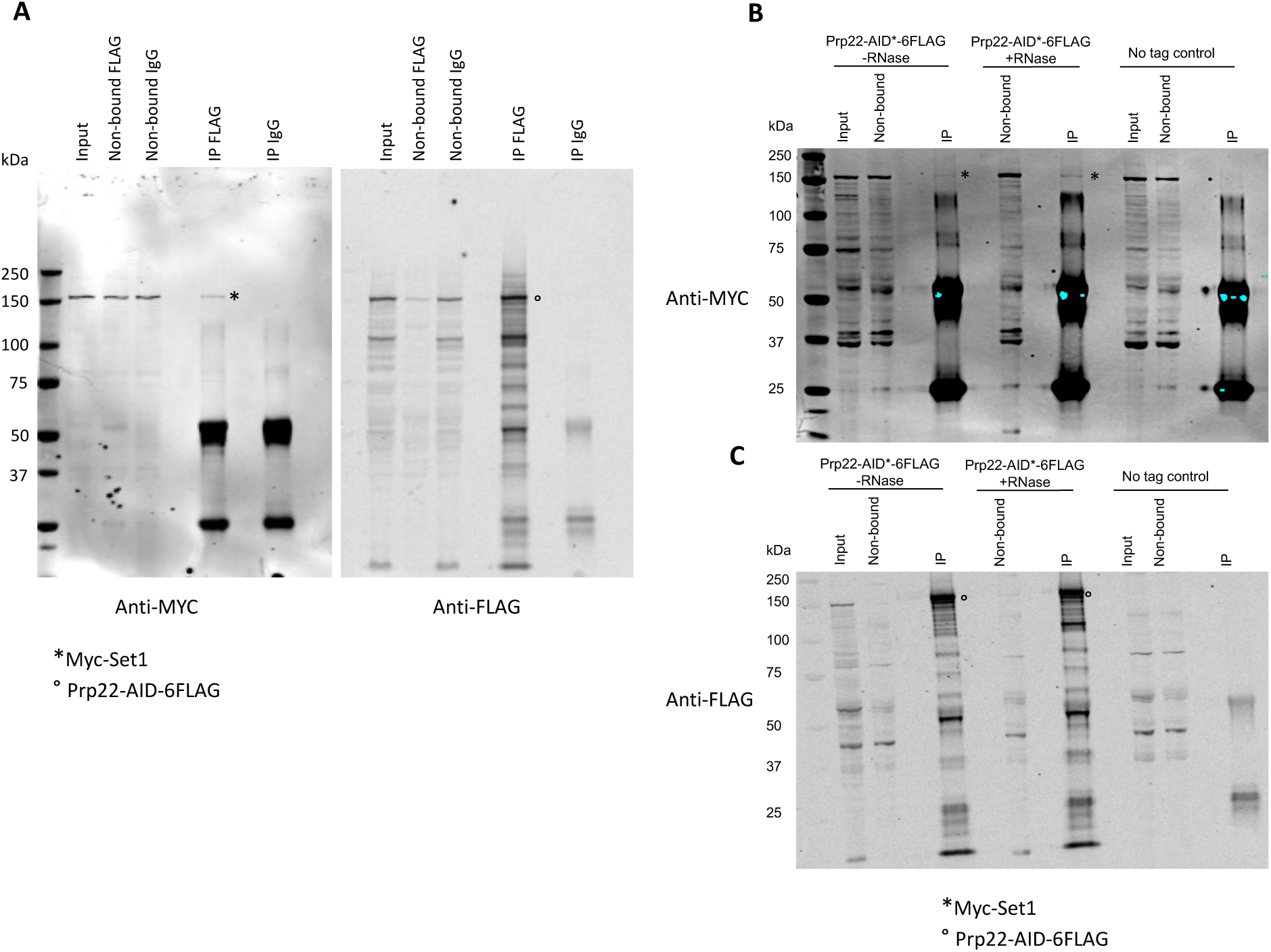
Set1 is co-precipitated with Prp22 *in vivo*. **A**) Co-immunoprecipitation experiments were performed in W303 expressing chromosomally encoded Myc-Set1 (Dehe et al. 2006) and Prp22AID-FLAG (Mendoza-Ochoa et al. 2019. Prp22-AID-6FLAG was pulled down using a FLAG antibody. IgG was used as a negative control. Input, non-bound and immunoprecipitation (IP) samples were loaded. The Western blot was probed either with an antibody against the MYC epitope tag (Left) or against the FLAG epitope tag (Right). Myc-Set1 and Prp22AID-FLAG are indicated by * and °, respectively. **B** and **C**) The same experiment was carried out in the absence or presence of RNase. A strain with 9MYC-Set1 and no flag-tagged Prp22 was used as a negative control. The Western blots were probed with anti-MYC and anti-FLAG respectively.

Finaly and somewhat surprisingly, we found that many Y2H interactors of Set1 1-754 were involved in tRNA nuclear transport, modification, and synthesis (Fig. 3). Swd2, which interacts with the Set1 N-terminus, exhibit a 2H interaction with a high degree of confidence with the cytoplasmic asparaginyl-tRNA synthetase Ded81 and the prolyl-tRNA synthetase Yhr020 while Sdc1interacted with the glycyl-tRNA synthase Grs1 and Bre2 with the glutamine tRNA synthetase Gln4. Of note, Trm1 (Liu *et al*, 1998) and Trm732 (Guy *et al*, 2012) are both directly or indirectly involved in tRNA methylation and form part of what has been defined as the cell’s global methytransferome (Giaever *et al*, 2019).

### SET1C Y2H interactors are involved in several aspects of DNA transactions

As mentioned in the introduction, it has been shown that SET1C, and in particular Spp1, regulate the selection of meiotic breaks, the progression of the replication fork, DNA repair, chromosome segregation, and transposition of Ty elements (Deshpande & Bryk, 2023). We have classified all Set1 and subunit interactors according to these SET1C roles (Fig. S4). We found a number of putative interactors involved in meiosis. In particular, Spp1 not only interacts with Mer2, but may also associate with the meiosis specific DNA helicase Mer3 (Nakagawa & Kolodner, 2002), which itself may interact also with Swd1 (Fig. S4). We further found Set1 candidate interactions with the kinetochore proteins Spc25 and Cbf2 suggesting that Set1 could be transiently localized at the spindle pole body. Regarding the biology of retrotransposons, we previously reported that Set1 binds post-transcriptionally to Ty1 mRNA and represses Ty1 mobility (Luciano *et al*, 2017). Interestingly, the two-hybrid screens reveal that Set1 1-754 interacted with Gag capsid-like proteins of Ty1 (Fig. S4) raising the possibility that Set1 binding to Ty1 mRNA is linked to the interaction of Set1 1-754 with Gag. Concerning the role of Set1 and subunits in DNA replication and repair, we report multiple potential interactions (Fig. S4). In particular, Swd2, Swd1, and Spp1 exhibited a high confidence 2H interaction with Orc6, Nrm1, and Mcm2, respectively (Table S2 and Fig. S4). Orc2 was previously described to interact physically with Spp1 (Kan *et al*, 2008). We recently reported that Spp1 is recruited at replication forks stalled at the Tus/Ter barrier independently of its interaction with Set1 (Ghaddar *et al*, 2023). Interestingly, in the Y2H screen Spp1 interacted with the extreme C-terminal region of Mcm2 (791-867) (Fig. S5A) that corresponds to a non-conserved accessible alpha-helix within the MCM complex (Li *et al*, 2015b). We fused Spp1 and Mcm2 to GST and performed GST pull-down experiments. We confirmed *in vitro* in both directions a weak interaction between Spp1 and Mcm2 (Fig. S5B). Whether Spp1 is recruited by Mcm2 at stalled replication fork remains to be determined.

Screening with *SWD1* yielded a high number of clones, fourteen of which contained fragments of the *NRM1/YNR009w* gene, which encodes a basic protein of 249 amino-acids (Table S2). Nrm1 inactivates MBF, a major regulator of the G1/S transcription (de Bruin *et al*, 2006). During replication stress, Nrm1 phosphorylation by the checkpoint kinase prevents its binding to MBF target promoters leading to the activation of G1/S transcription (Travesa *et al*, 2012). An exciting idea is that Swd1 recruits Nrm1 to stalled forks by promoting its phosphorylation by Rad53. Swd1 would play a role in linking replication stress and transcriptional regulation via Nrm1. Of note, Nrm1 was identified as a gene required for the cell-cycle pattern of H3K79me2 during early S phase (Schulze *et al*, 2009). Interestingly, Nrm1 exhibit a H3K4-like domain and raised our attention to other yeast proteins that carry such sequences. We used the scansite search algorithm (http://scansite.mit.edu) to systematically identify sequence motifs that are related to the SET1C modification site in histone H3. Search parameters included four to six identical residues of the ARTKQT sequence that is found at the H3K4 modification site. In addition, we did allow for biochemically equivalent amino acid changes. Fig. S6 shows eight candidate proteins based on their function and the sequence context of the H3K4-like domain. Some of these gene products are involved in cellular pathways that functionally overlap with SET1C, e.g. transcription (Not5), rDNA silencing (Irs4) and cell-cycle control (Dbf2 and Dbf20), increasing the possibility of occurring physical and functional interactions.

Finally, Y2H screening indicated that Set1 and its subunits exhibit 2H-interactions with a number of proteins involved in protein SUMOylation (Fig. S4). The proteins were either involved in SUMO conjugation or SUMO-dependent degradation. Remarkably, the C-terminus of Nis1 (360-407) that contains a potential SUMO-binding site (Hannich *et al*, 2005) was identified a high-confidence interactor of Spp1, Shg1, and Sdc1 (Fig. 1). Nis1 is localized at the bud neck (Iwase & Toh-e, 2001) at the vicinity of the septin collar containing several highly SUMOylated proteins (Shs1, Cdc11) (Wykoff & O’Shea, 2005) and has been implicated in preventing bud recovery at the site of division (Meitinger *et al*, 2014). We thus sought to confirm biochemically the interaction of Nis1 with the three SET1C subunits. We fused the GST to Spp1, Sdc1, and Sgh1 to perform pull-down experiments with *in vitro* translated Nis1. We confirmed the interaction of Nis1 with Spp1, and Sdc1, but not with Sgh1 (Fig. S7). However, mass spectrometry analyses on TAP-Nis1 did not reveal the presence of SET1C subunits (Table S3) suggesting that interaction between Nis1 and Spp1/Sdc1 might be transient. The relevance of the Nis1 and other putative interactors remains unclear, especially as many of those proteins are not known to be located in the nucleus. Interestingly, Nis1 has been reported to shuttle from the bud neck to the nucleus when overexpressed (Perez & Thorner, 2019). In Fig. S8, we have identified a number of interactors, including Sdc1 and Spp1, showing changes in localization following hypoxia (Henke *et al*, 2011). Of note, one of the two human SET1C homolog SET1B has a cytoplasmic location with functions unrelated to H3K4 methylation (Wang *et al*, 2017). Combined, these observations raise the question of the minor or transient localization of Set1 and its subunits outside the nucleus, or conversely of the transient localization of interactors within the nucleus.

### Set1 is SUMOylated

Global analyses of SUMOylated proteins in fission yeast revealed that Set1 and Spf1 (Spp1) were SUMOylated (Shin *et al*, 2005; Nie *et al*, 2015). As Set1 had two-hybrid interactions with Slx5 and Wss1, both of which act on SUMOylated proteins as they undergo protein degradation (Mullen *et al*, 2010), we tested whether Set1 can be SUMOylated. Cells expressing Myc-Set1 (Dehe *et al*, 2006) or transformed with pPB66-SET1 expressing GBD-Set1-FL (GAL4 binding domain) were transformed with a plasmid encoding His_6_-SUMO or a control plasmid. Proteins were purified on Ni-NTA agarose beads and detected by Western blot either using anti-MYC or anti-GAL4 antibodies. The results revealed that Set1 can be mono-SUMOylated (Fig. 5A) or di-SUMOylated (Fig. 5B). This difference may be due to the fact that GBD-SET1-FL is under the control of the ADH promoter in plasmid pGB66 and thus overexpressed.

**Figure 5.**
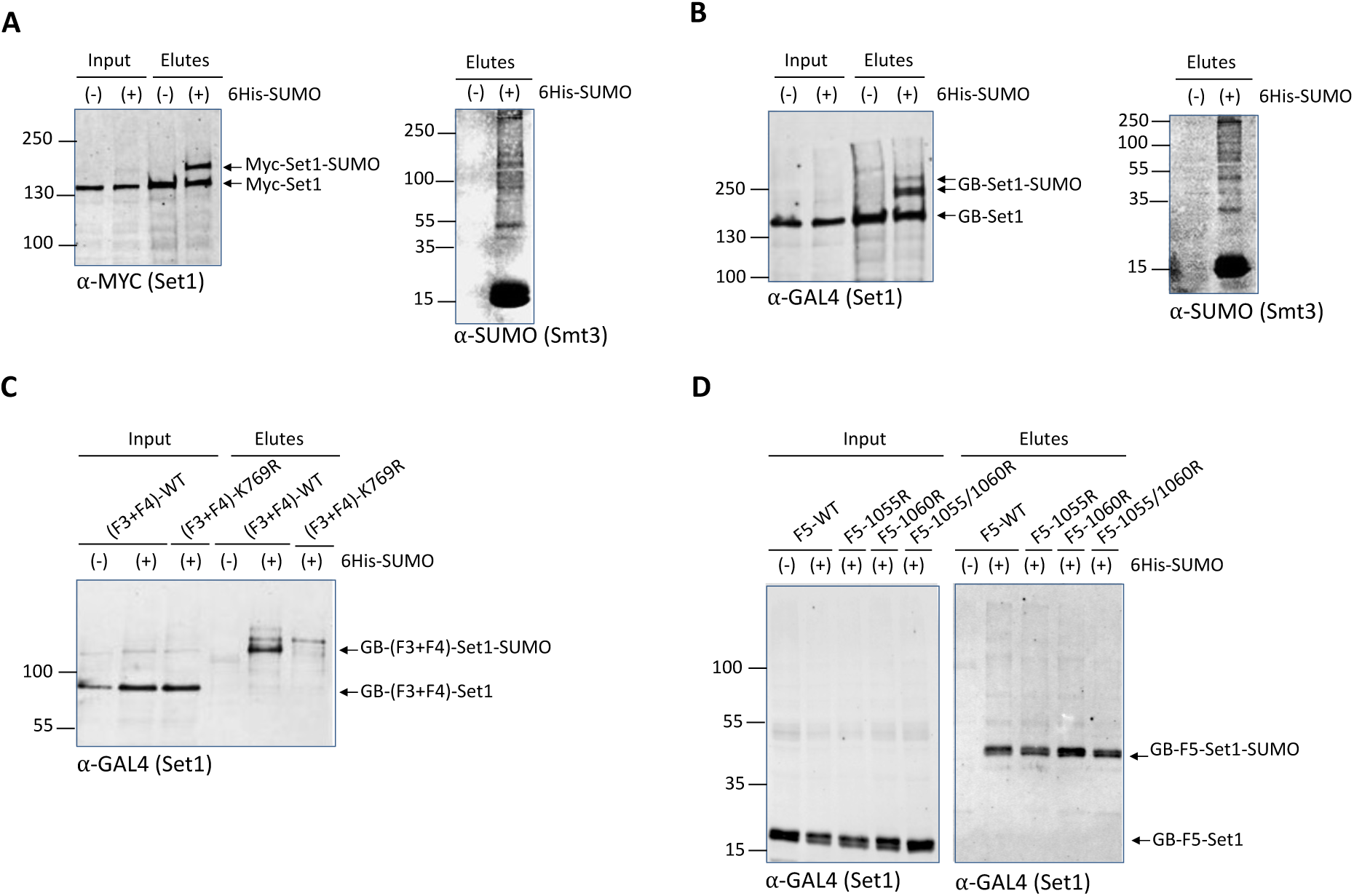
Set1 is SUMOylated. 6His-SUMO–conjugated proteins were purified from cells transformed (+) or not transformed (−) with a plasmid encoding 6His-SUMO under control of the *CUP1* promoter. Cell lysates (Input) and Ni-purified material (Elutes) were analyzed by Western blotting with an anti-MYC antibody (**A**) or and anti-GAL4 antibody (**B-D**). Analysis of 6His-SUMO-conjugated forms of (**A**) genomically MYC-tagged Set1 or (**B**) GB-Set1 transformed cells was performed (left panels), in both the cases SUMO expression and efficiency of purification were controlled using an anti-SUMO antibody (right panels). **C**) SUMOylation analysis of Set1 fragment F3+F4 (aa. 351-956) WT and the K769R mutant. **D**) SUMOylation analysis of Set1 fragment F5 (aa. 956-1080) WT, single mutants K1055R and K1060R, and the double mutant K1055R/K1060R mutant. Unmodified MYC-Set1 and GAL4-Set1 in both the (-) and (+) His-SUMO eluates are most likely due to the stickiness of unmodified Set1 to the beads.

We sought to refine the SUMOylated Set1 region. We transformed the plasmid encoding His6-SUMO and the control plasmid into cells expressing Set1 fragments F1, F2, F3, F4, F3+F4, and F5 (Fig. S9A). We expressed F3+F4 instead of the isolated F3 and F4 fragments in order to preserve the K769-centered motif predicted to be highly SUMOylated (Fig. S9B). F1, F2, F3, and F4 were not SUMOylated (not shown) while the F3+F4 fragment was clearly SUMOylated (Fig. 5C). In order to identify whether K769 is important for F3+F4 SUMOylation, we mutated K769 to R. We show that introduction of K769R into F3+F4 abolishes its SUMOylation (Fig. 5C), indicating that K769 is likely to be the SUMOylated lysine within the F3+F4 fragment. We then found that F5 was also SUMOylated (Fig. 5D). In this case, the substitution K1055R, K1060R, or the double substitution, does not affect the SUMOylation of F5 indicating that this motif also predicted to be SUMOylated with a high score is not the SUMOylated motif (Fig. 5D). Taken together, these experiments indicate that Set1 can be SUMOylated in the N-SET and SET domains, in the interaction region of Spp1 and Swd1-Swd2-Bre2-Sdc1, respectively. This raises the possibility that SUMOylation regulates the interaction of Set1 with its subunits, in particular with Spp1, which interacts dynamically with SET1C (Ghaddar *et al*, 2023; D’Urso *et al*, 2016; Karányi *et al*, 2018; Serra-Cardona *et al*, 2022). Interestingly, in mammals, the SUMO peptidase SENP3 (the ortholog of Ulp1 in budding yeast) interacts with MLL1 and MLL2, catalyzing the deSUMOylation of RbBP5 (Swd1). This process regulates the association of specific subunits of MLL1/MLL2, such as menin and Ash2L (Bre2), with the DLX3 gene, which plays a role in osteogenic differentiation (Nayak *et al*, 2014).

### SET1C Y2H interactors regulate metabolism and stress responses

Set1 was reported to regulate ergosterol levels (South *et al*, 2013). We identified Erg9, Ugt51, Vhr1, Vhr2 and Ste20, all involved in ergosterol metabolism (Daicho *et al*, 2020; Lees *et al*, 1995; Warnecke & Heinz, 1994). Erg9 and Ste20 were candidate interactors with Set1 while Ugt51 and Vhr1/Vhr2 were high confidence 2H-interactors of Shg1 and Sdc1, respectively (Fig. S10). Along the same line, the various Y2H screens revealed numerous genes involved in phosphatidyl inositol metabolism as potential interactors of Set1 and its subunits. In particular Vip1 is a high-confidence 2H-interactor of Set1 754-1081 (Fig. 1B). Vip1 is a bifunctional inositol pyrophosphate kinase and phosphatase that regulates IP7 levels in the inositol pyrophosphate (PP-IP) synthesis pathway (Lee *et al*, 2007; Mulugu *et al*, 2007). Interestingly, Vip1 was reported to regulate the environmental stress response (ESR) through IP7 that activates the HDAC Rdp3 (Worley *et al*, 2013) suggesting potential new avenues to explain ESR regulation by Set1 (Weiner *et al*, 2012). Along the same line, Swd1 and Swd3 were identified in a screen aimed to identify genes that negatively regulate the PHO pathway in a Vip1-dependent manner (Choi *et al*, 2017). How the putative interaction between Set1 754-1081, which also interacts with Pho23 subunit of the Rpd3L complex, and Vip1 fits into these processes remains to be discovered. Of note, Vip1 has been shown in *Arabidopsis thaliana* to change localization upon hypo-osmotic stress from the cytosol to the nucleus (Takeo & Ito, 2017). On their side, Y2H-interactors identified as stress-responsive genes interact either with Set1 1-754 or Swd2 (Fig. S10). Swd2 interacted with high confidence with Nar1, an essential Fe/S protein required for the assembly of cytosolic Fe/S proteins (Balk *et al*, 2004) and the calcineurin phosphatase Cmp2 activated in response to ER stress (Mizuno *et al*, 2018). In contrast to the stress genes mentioned above, many interactors involved in glucose metabolism (repression) interact only with the Set1 1-1081 or with members of the nSET module (Fig. S10).

### Reconstituted Set1C methylates the Snf2 RG motif within the AT-hook *in vitro*

Our screens indicate that Snf2 (residues 1430-1499) was potentially interacting with Set1 1-754 (Table S2). To validate this interaction we obtained recombinant protein where GST was fused to the C-terminal domain of Snf2 (Snf2C), the AT-hook (Snf2C-AT-hook) and the Bromo domain (Snf2C-Bromo) (Kim *et al*, 2010) (Fig. 6A). We performed pull-down assays to test the interaction of the GST-fusion proteins with reconstituted SET1C or with the SET1C762 complex in which Set1 has a N-terminal truncation of the first 761 residues, respectively (See METHODS) (Fig. S11 A-E). We found that SET1C as well as the SET1C762 complex were both associated with Snf2C and Snf2C-AT-hook but not to Snf2C-Bromo (Fig. 6 B, C), indicating that AT-hook is a key region for interaction with SET1C. We then delineated the minimal AT-hook region required for interaction with SET1C and found that residues 1461-1547 are essential for binding to SET1C (Fig. 6 D-E). Collectively, these results indicate that SET1C interacts with Snf2C-AT-hook region suggesting that the Set1 and Snf2 complexes cooperate to modify chromatin structure.

**Figure 6.**
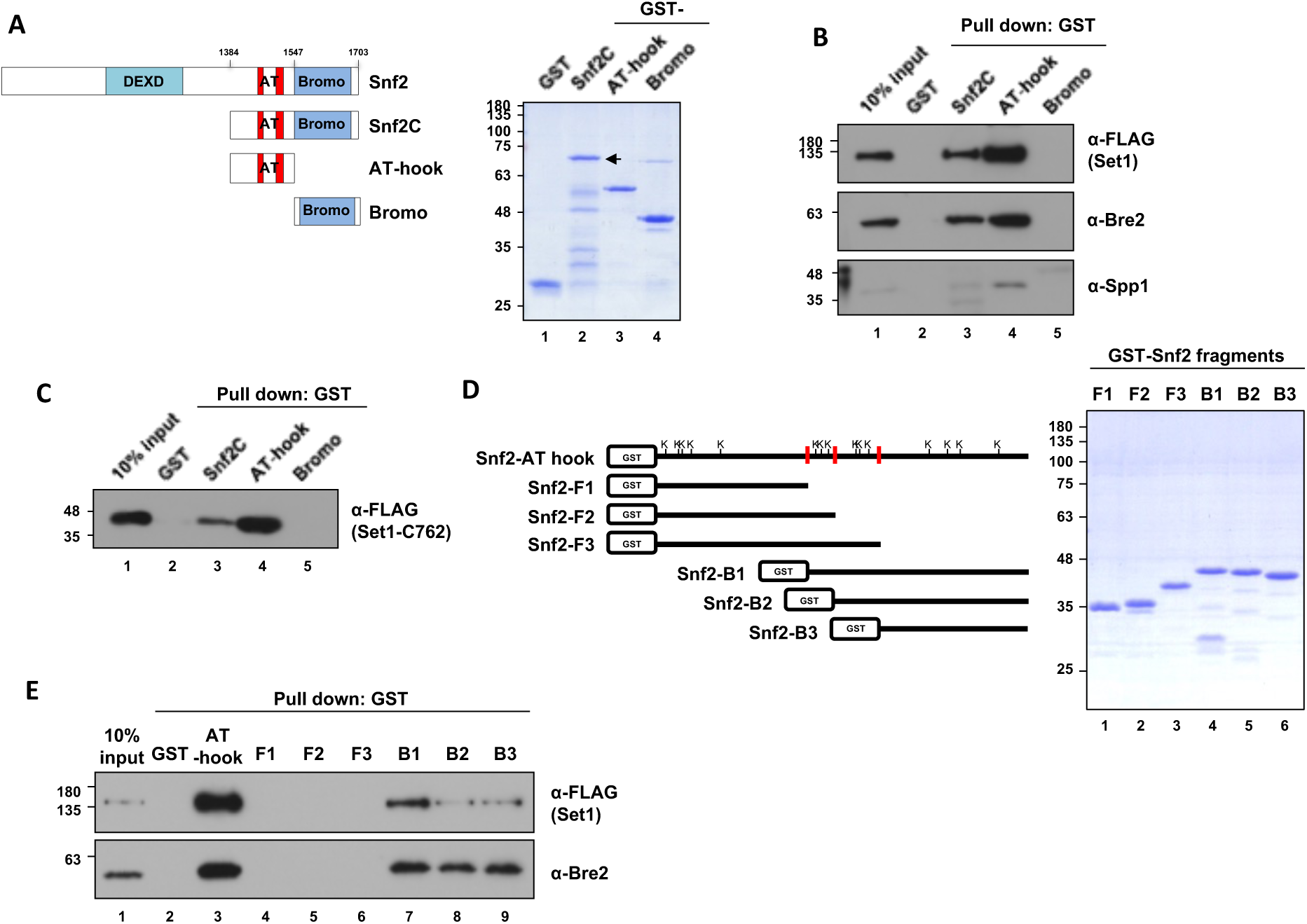
SET1C interacts *in vitro* with Snf2C-AT-hook. **A**) A schematic diagram depicting the domains of Snf2 and the Snf2 fragments used in this study, along with the SDS-PAGE/Coomassie staining of the purified GST-tagged Snf2 fragments. **B** and **C**) GST pull-down assay using purified GST-tagged Snf2 fragments. The purified SET1C (**B**) or SET1C-C762 complex (**C**) was mixed with GST-tagged Snf2 fragments, followed by GST pull-down, and the bound proteins were analyzed by immunoblotting. **D**) A schematic diagram illustrating Snf2 fragments with a more detailed breakdown of the AT-hook domain, along with the SDS-PAGE/Coomassie Blue staining of the purified Snf2 fragments. The lysines present in the AT-hook are represented by the letter K. **E**) GST pull-down assay using purified GST-tagged Snf2 fragments and SET1C.

We have shown above that SET1C interacts with Snf2C-AT-hook region (Fig. 6). We thus tested whether the Snf2C-AT-hook could be methylated by Set1C. Set1C and truncated Set1C were reconstituted and affinity purified from Sf9 cells coinfected with baculoviruses that express FLAG-Set1 (or truncated Set1) and the seven other untagged subunits (Kim *et al*, 2013). We incubated reconstituted Set1C with the Snf2C, AT-hook and Bromo fragments in the presence of radioactive S-adenosylmethionine (^3^H-SAM). The results indicate that Set1C methylates *in vitro* purified Snf2C and Snf2C-AT-hook, but not Bromo (Fig. 7A, B) and that Snf2C-AT-hook methylation requires Set1 FL (Fig. 7C). Collectively these results show that the Snf2C-AT-hook (1384-1547) interacts and is methylated by Set1 FL *in vitro*. Lys 1494 (K1494) and Lys 1498 (K1498) located between the AT-hook domains of Snf2 were previously shown to be acetylated by Gcn5 (Kim *et al*, 2010). We individually mutated all the Lys of the Snf2-AT-hook into Arg and tested the methylation of the mutated fragments. None of the substitutions abolished or decreased the methylation of the AT-hook indicating that he AT-hook could be methylated on multiple sites or on other type of residues (Fig. S12). We thus sought to identify the minimal region within the Snf2-AT-hook that is methylated by Set1C *in vitro*. The Snf2-AT-hook domain was divided in 6 regions that were fused to the GST (Fig. 7D). *In vitro* methylation assays indicated that the Snf2-B1, B2, B3 were methylated by the reconstituted Set1C (Fig. 7E). As the Snf2-B3 fragment that contains the last four lysines (K1488, K1494, K1498, K1526) of Snf2-AT-hook domain was still methylated, it suggested that one of several of these lysines are methylated by Set1C. We then examined which of these 4 lysine residues are methylated by Set1C *in vitro*. To our surprise, mutating all the lysine residues of the Snf3-B3 fragment did not abolish the methylation of Snf3-B3 (Fig. 7F). These results led us to think that Set1C could methylate arginine residues, in particular those contained in the RGG motif of the Snf2-B3 fragment. We thus deleted the RG repeats of Snf2-B3 (Fig. 7G) and purified the *Snf2-B3ΔRG*. We found that deletion of the Snf2-B3 RG repeats abolished the interaction between Snf2-B3 and Set1 and methylation of Snf2-B3 (Fig. 7H, I). The fact that both, interaction and methylation are lost upon deletion of the RG motif argues in favor that Set1C, and not a potential contaminant from the reconstituted Set1C, is responsible for the methylation of the Snf2-B3 RG repeats. Individual deletion of pairs of arginine residues in the RG motif did not suppress methylation of the Snf2-B3 fragment, suggesting flexibility in Set1C’s ability to methylate Snf2-B3 (Fig. S13).

**Figure 7.**
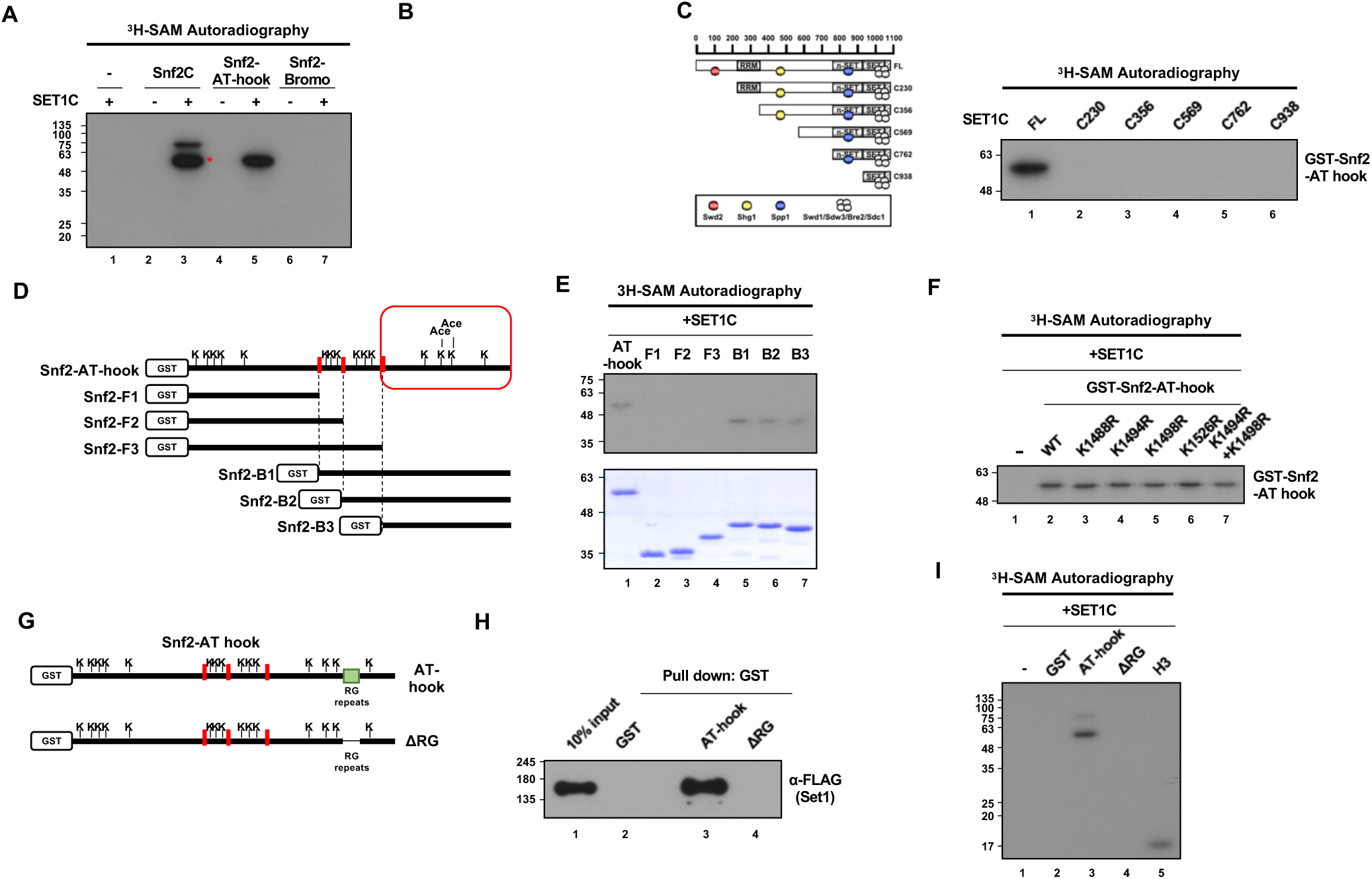
Snf2 is methylated in tandem AT-hook domain by reconstituted Set1C. **A**) *In vitro* methyltransferase assay using purified SET1C and Snf2 fragments. ^3^H-SAM was used as a methyl-donor and methylated proteins were detected by autoradiography. The band marked with a red star is a degradation product of Snf2C. **B**) *In vitro* methyltransferase assay using SET1C and two Snf2-AT-hook fragments with two different tags. **C**) Schematic diagram of N-terminal truncated SET1 complexes (left) and *in vitro* methyltransferase assay with GST-Snf2-AT-hook and truncated SET1 complexes. **D**) Schematic diagram showing the positions of all lysines in Snf2-AT-hook and the further cleaved fragments of Snf2-AT-hook. The red box indicates the two lysines that are acetylated by Gcn5. **E**) Coomassie staining of purified Snf2 fragments (lower) and an *in vitro* methyltransferase assay using these fragments with SET1C (upper). **F**) *In vitro* methyltransferase assay by SET1C when each of the four lysines in the C-terminal region of the Snf2-AT-hook is substituted with arginine or when both lysines known to be acetylated by Gcn5 are substituted. **G**) A schematic diagram showing the position of the RG-repeat region and the design of Snf2-AT-hook with RG-repeat truncation. H and I) GST pull-down assay (**H**) and *in vitro* methyltransferase assay (**I**) using purified SET1C and GST-Snf2-AT-hook with or without RG-repeats.

To further confirm that Snf2-B3 is methylated on arginine, we incubated a mutant Snf2-B3 (in which lysines were mutated in alanine to limit protease digestion prior mass spectroscopy analysis) with reconstituted Set1C and SAM and further purified the Snf2-B3 mutant on a Ni-NTA resin (Fig. 8A-D). We then analyzed by mass spectroscopy arginine methylation in Snf2-B3. We found that R1490, R1501, R1505, R1507 and R1517 were mono- and di-methylated while R1509, R1513, and R1519 were monomethylated (Fig. 8E). We thus confirmed that reconstituted Set1C has the ability to methylate in vitro multiple arginines in the Snf2B3 region.

**Figure 8.**
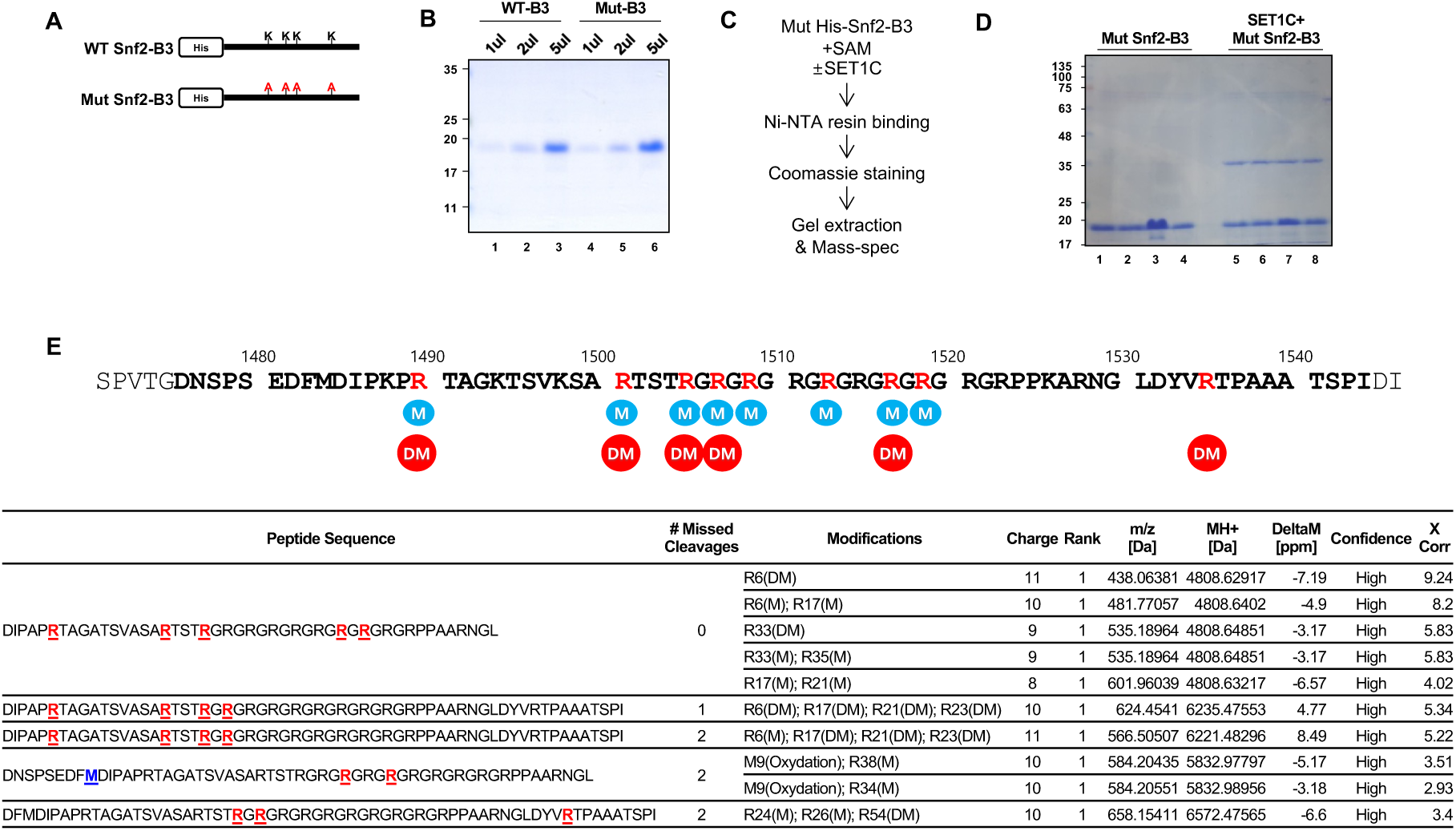
The arginines in the RG-repeat of Snf2 are methylated by reconstituted SET1C. **A**) A diagram showing the WT Snf2-B3 fragment and the Mut Snf2-B3 with all four lysines substituted with alanine. **B**) Coomassie staining of purified WT and Mut Snf2-B3. **C**) Mass-spectrometry experiment design to identify the methylation sites of Snf2-B3. **D**) Coomassie staining of Mut Snf2-B3 after methylation reaction and an additional purification step using Ni-NTA. The band corresponding to Mut Snf2-B3 (∼18 KDa) was excised and used for mass spectrometry analysis. The ∼37 KDa band observed in lanes 5–8 appears to be a SET1C subunit that binds non-specifically to Ni-NTA, likely SWD2 based on its size. **E**) Mass spectrometry analysis result of Snf2-B3 methylation sites revealed that multiple arginines in the RG-repeat were methylated. Amino acid sequence of Snf2 highlighting the arginines methylated by SET1. The sequence of the B3 fragment is shown in bold, and the arginines methylated by SET1 are marked in red. Methylation and demethylation are denoted as M and DM, respectively.

### Snf2 is methylated *in vivo* on its ARTSTRGR AT-hook motif in a Set1-dependent way

Our results show that an activity associated with reconstituted Set1C is capable of methylating arginines in the vicinity and in the RG repeats of Snf2C-AT-hook. The fact that only Set1FL is capable of methylating the AT-hook is in favor of this activity being associated with Set1C. Nevertheless, one cannot formally rule out that methylation the AT-hook is due to a contaminant from insect cells from which Set1C has been reconstituted.

We thus investigated the interaction between Snf2 and Set1 *in vivo* and whether Snf2 is methylated by Set1C. We found that Set1 was coimmunoprecipitated with Snf2 tagged with a C-terminal Myc epitope (Fig. 9A). In contrast to what we observed *in vitro*, deletion of the RG motif of Snf2 did not affect the interaction between Snf2 and Set1, which is consistent with the fact that the SID of Snf2 is located upstream of the RG repeats (Fig. 9A).

**Figure 9.**
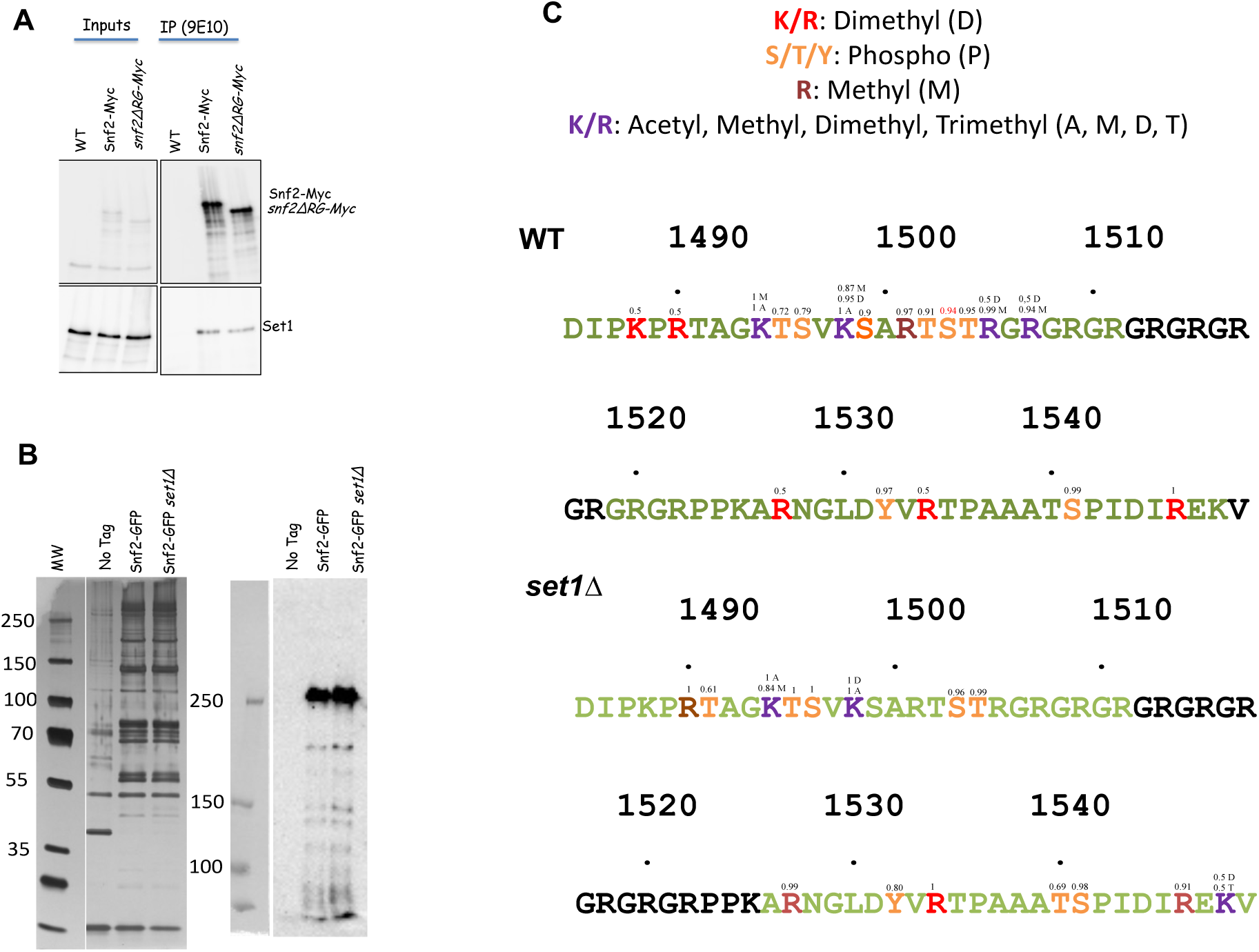
The arginines of the motif ARTSTRGR within the AT hook are methylated *in vivo* in a Set1-dependent way. **A**) Set1 interacts *in vivo* with Snf2 and Snf2-ΔRG. Myc-tagged Snf2 and Snf2ΔRG were immunoprecipitated with 9E10 Myc antibodies (see Methods) and revealed with either 9E10 (Upper panel) or Set1 antibodies (lower panel). **B**) Snf2C complex was purified from WT and *set1Δ* strains, separated on a 4-12% Bis-Tris Gel and Silver Stained (*Left*); the presence of Snf2-GFP is detected by Western-blotting with anti-GFP antibody (*Right*). The area corresponding to Snf2-GFP was excised from the gel and used for mass spectrometry analysis. Peptides flanking the RG repeats (1485-1549) with their PTM are shown in Fig. S15. **C**) Panel C show a focus on the amino sequence flanking the RG repeats. The positions of residues from D_1485_ to V_1549_ are indicated on the figure. Peptides identified after digestion of Snf2-GFP are indicated in color with their identified PTM indicated by the color code shown in the top of the panel. The small numbers above the amino acids indicate the probability of localisation according to the MS2 peaks. It should be noted that for wild-type K1488 and R1490, a peptide with dimethylation is detected, but no discriminating MS2 peak allows us to conclude whether K or R are dimethylated. This is also the case for dimethylation on R1528 and R1535. The results presented represent the observed PTMs from two independent experiments, each containing 3 replicates (Table S6).

To further investigate Snf2 methylation by Set1C *in vivo*, we purified Snf2-GFP and its subunits with GFP nanobodies in the presence or absence of Set1 (Fig. 9B). The protein composition of the complex was determined by mass spectrometry in the presence of absence of Set1 (Fig. 9B, Table S5). We recovered in both cases all the known components of the Snf2 complex (Snf2, Swi1, Swi3, Snf5, Snf12, Swp82, Arp9, Arp7, Snf6, Rtt102, Snf11, Taf14) (Table S5, Fig. S14). Interestingly, we found that the arginine methyltransferase Rmt2 was found specifically enriched with the Snf2-GFP complex in *set1Δ* cells (Fig. S14).

We then excised the Snf2-GFP gel band and subjected it to mass spectrometry. We examine the post-translational modifications (PTM) in the Snf2 (1485-1549) region in either WT or *set1Δ* strains (Table S6). We were unable to purify a peptide containing the RG repeats but we could identify the peptides flanking the RG motif (Fig. 9C). Fig. 9C represents the combined results of two independent experiments. In WT cells, we found that the region between K1494 and R1505 was reproducibly subjected to multiple PTM (Fig. 10C, upper panel). K1494 and K1498 that were previously found acetylated by Gcn5 (Kim *et al*, 2010) were found acetylated, however both lysines were also found to be methylated (Fig. 9C). Of note H3K4 that is methylated by Set1C has been reported to be also acetylated by Gcn5 (Guillemette *et al*, 2011). T1995, S1996, S1999, T1502, S1503, and T1504 were found phosphorylated. Interestingly, R1501 was found mono-methylated while R1505 and R1507 were found di-methylated in the two independent Snf2-GFP purifications. These 3 arginines belong to a ARTSTRGR motif that lies in the Snf2-B3 fragment. They were also found to be methylated *in vitro* by the reconstituted Set1C (Fig. 8E). We then examine the posttranslational modifications of the peptides flanking the RG motif in the *set1Δ* strain (Fig. 9C, lower panel). Phosphorylation of serines and threonines was not modified in the *set1Δ* strain except for T1502. In the *set1Δ* strain, K1494 and K1498 remained methylated indicating that methylation of both lysines does not depend on Set1C. Strikingly, deleting *SET1* abolished the mono-methylation of R1501 and the di-methylation of R1505, R1507 (Fig. 9C, lower panel). The results thus show that SET1C is required for the methylation of the three arginines of the ARTSTRGR motif, in agreement with the *in vitro* results.

## DISCUSSION

We have exploited the power of Y2H screening technology to establish an extended interaction network for the histone H3K4 methylase complex SET1C. Comprehensive data integration unveiled many potential functions of either the whole complex or individual subunits. This study is partly validated by the fact that 2H-interactors were found in known functions of SET1C, or putative functions for which the mechanism is unknown. We have also provided a limited number of validations by confirming the interaction through biochemical approaches. It seems likely that some proteins may interact transiently with Set1 and its subunits, or under specific conditions of stress or nutrient limitations, explaining why these interactors have not previously been found associated with SET1C in biochemical approaches. The most illustrative case for this is the high-affinity interaction between Spp1 and Mer2 that takes place during the first meiotic division, which we had identified from the Spp1 screen and which has been extensively characterized (Acquaviva *et al*, 2013b). Nevertheless, considering the multitude of interactions identified, it seems unlikely that they all represent direct contacts. The list shown in Table S2 may contain multiple false positives and further work will be required to determine whether these interactions occur in a physiological context and what their functional significance may be. Another caveat is that Y2H interactions can be mediated by endogenous proteins that bridge the interactions.

However, we anticipate that this systematic screen will be an invaluable resource for further investigation of the role of SET1C in known and novel processes revealed in this study. Processes discussed here include Set1’s export/import mechanism through the nucleus, interaction with RGG proteins some of which are targeted by arginine methyltransferase, cooperation of SET1C with chromatin remodeling factors in particular Snf2C, interaction with splicing factors notably with Prp8 and Prp22, coupling replication with histone deposition (Mcm2/Spp1), SET1C plasticity regulated by SUMOylation, and Ty element transposition. A recurrent question related to the identification of potential SET1C interactors is whether they could be methylated by SET1C.

We show that Set1C interacts both *in vitro* and *in vivo* with Snf2. Strikingly, reconstituted Set1C from insect cells is able to methylate *in vitro* the three arginines located within the A**R**TST**R**G**R** motif of the AT-hook of Snf2. The *in vivo* results confirm that Set1 is required for the methylation of these arginines within the A**R**TST**R**G**R** motif. It is interesting to note that this motif resembles the A**R**TKQTA**R** motif of the N-terminal H3. In yeast, H3R2 is mono- and di-methylated and its asymmetric di-methylation prevents H3K4me3 (Kirmizis *et al*, 2009, 2007). In contrast symmetric di-methylation of H3R2 correlates with H3K4me3 and requires Set1 *in vivo* but not the classical arginine methyltransferases: Hmt1, Hmt2, Rkm2, Rkm3, Rkm4, Hsl7, and Efm1 (Li *et al*, 2015a; Yuan *et al*, 2012). It has been proposed that Set1C could be responsible for both H3R2me2s and H3K4me3 or that H3R2me2s deposited by a yet unidentified arginine methyltransferase requires the prior deposition of H3K4me3 (Yuan *et al*, 2012).

Interestingly, previous tandem affinity protein purification of the mono- and asymmetric arginine methyltransferase Hmt1 (also termed Rmt1) identified 108 proteins associated with it (Jackson *et al*, 2012), a number of which were shown to be methylated. Snf2 was found to be copurified with Hmt1 *in vivo* and methylated *in vitro* by Hmt1 (Jackson *et al*, 2012) making Hmt1 a potential candidate for methylating Snf2. *In vitro*, purified Hmt1 catalyzes mono-methylation and asymmetric di-methylation of H3R2, however, deleting *HMT1* does not affect the asymmetric di-methylation of H3R2 *in vivo*, which suggests that several methyltransferases could act redundantly on H3R2 (Li *et al*, 2015a). In mammals, H3R2 and H3R8 are asymmetrically di-methylated by PRMT6 (Hamey *et al*, 2021; Guccione *et al*, 2007; Hyllus *et al*, 2007). It has been also reported that PRMT5 can be found associated with hSWI/SNF and has the ability to methylate H3R8 (Pal *et al*, 2004). We did not identify Rmt1 in our purification but found that the arginine mono-methyl transferase Rmt2 enriched with Snf2-GFP in *set1Δ* cells. Rmt2 is a type IV methyl transferase that was reported to methylate the ribosomal protein L12 (Low & Wilkins, 2012; Chern *et al*, 2002). We could envision the possibility the possibility that Set1C could cooperate with distinct protein arginine methyl transferases to promote the mono-methylation of R1501 and the di-methylation R1505 and R1507 within the A**R**TST**R**G**R** motif. However, it was previously reported that the combined inactivation of Rmt1, Rmt2 and Hsl7 did not affect the mono-methylation of H3R2, weakening this hypothesis (Kirmizis *et al*, 2009). It is possible that in yeast there is a redundancy of enzymes capable of methylating the Snf2 A**R**TST**R**G**R** motif, but that for each of them their activity on this motif depends on the presence of Set1. Alternatively, Set1C could directly methylate the A**R**TST**R**G**R** motif as discussed for the A**R**TKQTA**R** H3 motif (Yuan *et al*, 2012). The fact that Set1C interacts with Snf2, Gbp2, Nop1, Nab2, Dbb1 (Fig. 2), all of which have RG motifs and are mostly Hmt1 substrates, raises the possibility of a general interplay between methylarginine and methyllysine.

Interestingly, a very recent article shows that PRMT1 binds to the N-terminal region of MLL2 and methylates multiple arginine residues within its RGG/RG motifs (An *et al*, 2025).

## MATERIALS and METHODS

### Strain construction

All strains and plasmids used in this study are listed in Table S1. To obtain gene deletions and expression of tagged proteins, we amplified by PCR a disruption or a tagging cassette containing the appropriate marker as described (Janke *et al*, 2004).

### Yeast two-hybrid analysis

Yeast two-hybrid screening was performed by Hybrigenics Services, S.A.S., Evry, France. The coding sequence for Set1-FL, Set1 1-754, Set1 754-1081 and Bre2 were cloned into pB66 (GAL4-bait) as C-terminal fusions to the Gal4 DNA-binding domain while those of Swd1, Swd3, Sdc1, Spp1, and Shg1 were cloned into pB27 as a C-terminal fusion to LexA (LEXA-bait). *SWD2* was cloned into pB43 as a N-terminal fusion (SWD2-GAL4). Individual bait cloning was performed by using specific primers for PCR for every bait. Every PCR fragment subcloned as bait is entirely sequenced to avoid eventual mismatches in the coding sequence. The constructs were used as baits to screen a genomic *S. cerevisiae* library constructed into pP6 based on the pGADGH plasmid (Wilson *et al*, 1993). pB6, pB66 and pB43 are derived from the original pAS2Δ vector (Fromont-Racine *et al*, 1997). The library was submitted to very strict quality controls. Each protein is represented by several fragments (domains) and the library has been screened with a set of 6 reference baits before any other bait proteins are screened.

The Gal4 constructs were screened using a mating approach with YHGX13 (Y187 *ade2-101::loxP-kanMX-loxP, MATalpha*) and CG1945 (*MATa*) yeast strains. The LexA constructs were screened using a mating approach with YHGX13 (Y187 *ade2-101::loxP-kanMX-loxP, mat alpha) and L40ΔGal4* (*MATa*) yeast strains as previously described (Fromont-Racine *et al*, 1997). Positive clones were selected on a medium lacking tryptophan, leucine and histidine and supplemented with 3-aminotriazole (3AT) if necessary to handle bait auto-activation. The prey fragments of the positive clones were amplified by PCR and sequenced at their 5’ and 3’ junctions. The resulting sequences were used to identify the corresponding interacting proteins in the GenBank database (NCBI) using a fully automated procedure.

A confidence score (PBS, for Predicted Biological Score) was attributed to each interaction as previously described (Formstecher *et al*, 2005). The confidence score relies on two different levels of analysis. Firstly, a local score takes into account the redundancy and independency of prey fragments, as well as the distribution of reading frames and stop codons in overlapping fragments. Secondly, a global score takes into account the interactions found in all the screens performed at Hybrigenics using the same library. This global score represents the probability of an interaction being nonspecific. For practical use, the scores were divided into four categories, from A (highest confidence) to D (lowest confidence). A fifth category (E) specifically flags interactions involving highly connected prey domains previously found several times in screens performed on libraries derived from the same organism. Finally, several of these highly connected domains have been confirmed as false-positives of the technique. The PBS scores have been shown to positively correlate with the biological significance of interactions. When possible, the bait interacting domain of each prey is provided. The PBS scores have been shown to positively correlate with the biological significance of interactions (Rain *et al*, 2001). e-values for the interactions are available in the Hybrigenics database.

### Protein expression and protein interaction assays

Protein expression was done in *Escherichia coli* BL21 cells and purification of GST-fusion proteins were as described (Dichtl *et al*, 2002). MBP-fusion proteins were purified according to manufacturer’s instructions (New England BioLabs, Beverly, MA, USA). For His-tagged proteins expressed in bacteria, cDNAs were inserted into the pET28 vector (Novagen), expressed in *Escherichia coli*, and purified using Ni-NTA beads (Qiagen) following the previously described procedure (Kim & Roeder, 2011). GST pull-down assays were done as described (Dichtl *et al*, 2002).

A baculovirus expression system was used to express and reconstitute Set1 complexes containing either FLAG-Set1 or FLAG-Set1-C762. cDNAs were subcloned into pFASTBAC1, with or without an epitope tag, and baculoviruses were produced following the manufacturer’s instructions (Gibco-Invitrogen). Sf9 cells were infected with various combinations of baculoviruses, and the complexes were purified by affinity chromatography using M2 agarose, as previously described (Kim & Roeder, 2011).

### TAP-Nis1 affinity purification and mass spectrometry analysis

TAP-Nis1 and control cells (W303a) were grown in YPD, harvested in logarithmic phase (OD600 0.6-0.85) and cryo-lysed as previously described (Trahan & Oeffinger, 2022). Affinity purification was performed in RNP100 (20 mM HEPES-KOH pH 7.4, 100 mM NH_4_OAc, 0.5% Triton X-100, 0.1% Tween 20, 1:100 solution P, 1:5000 antifoam A, 100 mM NaCl) as described in (Trahan & Oeffinger, 2022). Following washes, the samples were on-bead trypsin digested in a volume of 50µl (f. c. 20 μg/mL trypsin (Sigma, proteomics grade) in 20 mM Tris-HCl (pH 8.0), 37 °C, 900 rpm, 16-20 h; stopped with 2% formic acid) and analyzed by tandem mass spectrometry as described in (Trahan & Oeffinger, 2022) using a 70-minute gradient on a LTQ Orbitrap Velos is a hybrid mass spectrometer (ThermoFisher Scientific) in Data-dependent mode. Data were processed with Thermo Excalibur to generate a raw file. Mascot search server (53 Version 2.3.02) was used with a parent tolerance of 10 ppm for precursor ions, 0.52Da for fragments, and only considering 1 possible missed cleavage as well as a mass change of +16 for methionine oxidations in the mass calculation. Data were searched against *S. cerevisiae* NCBI database and analyzed in Scaffold (version 3.6.4). The threshold and false discovery rates (FDR) were set to 80% and 0.37% respectively for peptides, and to 95% (1 peptide minimum) and 1.9% for respectively for proteins. Exclusive spectrum counts (ESC) were used for analysis and for each prey, the highest values obtained in the controls were removed from those of the samples during analysis.

### *In vitro* methylation

*In vitro* methylation reactions were done essentially as described (Roguev *et al*, 2001). 30 ml reactions typically contained 2 to 4 ml of partially purified SET1C complex, 2 to 4 mg substrate (either peptide, or core histone mixture, or recombinant H3) and 2 ml S-adenosyl (methyl-^3^H) methionine in MTA buffer (50 mM Tris 8.5, 20 mM KCl, 10 mM MgCl_2_, 250 mM sucrose). Reactions with core histones and recombinant H3 were resolved on 4-20% NuPAGE gels and subjected to fluorography. For Snf2 methylation assays, reaction mixtures containing purified SET1C (with 30 ng of the Bre2 subunit) and 200 ng of Snf2 fragments in 20 μl of reaction buffer (25 mM HEPES [pH 7.6], 50 mM KCl, 5 mM MgCl2, 0.1 mM EDTA, and 10% glycerol) were supplemented with 1 μCi of S-adenosyl (methyl-^3^H) methionine (PerkinElmer) and incubated at 30°C for 1 hour. Proteins were resolved by SDS-PAGE and subjected to fluorography. For fluorography, gels were fixed for 30 min in 40% methanol, 10% acetic acid, treated with EN^3^HANCE solution (Perkin Elmer) for 60 min, washed in cold dH_2_0 for 30 min, dried and exposed for five to fourteen days to photographic film.

### Mass spectrometry-based identification of Arg methylation in Snf2-B3

Mutant Snf2-B3 underwent a methylation reaction with SET1C and SAM, followed by purification via Ni-NTA affinity chromatography and separation by SDS-PAGE. The gel bands corresponding to the mutant Snf2-B3 were excised and subjected to in-gel digestion with AspN enzyme, followed by peptide extraction. The resulting peptide fractions were analyzed using an Easy-nLC 1200 coupled to an Orbitrap Fusion Lumos mass spectrometer (Thermo Fisher Scientific, MA, USA) at the Korea Basic Science Institute (Ochang). Peptides were separated on a C18 column using a 150-minute gradient, and data were acquired in data-dependent acquisition (DDA) mode. Full MS scans were acquired in the Orbitrap at a resolution of 60,000 over an m/z range of 350–2,000. Precursors for MS/MS analysis were selected for higher-energy collisional dissociation (HCD) fragmentation at a normalized collision energy of 30%. Raw MS data were processed using Proteome Discoverer with the SEQUEST search engine. The data were searched against the sequence of Snf2-B3 with a precursor mass tolerance of 10 ppm and a fragment mass tolerance of 0.02 Da. AspN was set as the digestion enzyme, allowing for up to two missed cleavages. Methylation, dimethylation, and methionine oxidation were specified as variable modifications. The false discovery rate (FDR) was set to 1% at the peptide level.

### Y2H interaction of Set1 fragments with selected preys

The Set1 fragments were amplified from the pB66-Set1-FL plasmid and cloned into the SfiI site of pB66. The selected preys (Snf2, Prp8, and Prp22) into p6 (Hybrigenics) were extracted from the screen. Plasmid pB66 contains the Gal4 DNA binding domain and the *TRP1* marker while pP6 expresses the Gal4 activating domain and the *LEU2* marker. TOTO cells were transformed with the different pB66-Set1-Fragments and the pP6-interactors and incubated 3 days at 30°C on SD-LEU-TRP. The transformant colonies were then streaked on SD-LEU-TRP-HIS containing either 5 mM or 20 mM of 3AT. To visualize interaction between Set1 fragments and the interactors, yeast cells were incubated 2 days at 30°C and cell growth was examined.

### AlphaFold model

AlphaFold models were generated as described in Abramson et al. (2024).

### CO-IP of Prp22-FLAG with Myc-Set1

Co-immunoprecipitation experiment was performed in W303 expressing chromosomally encoded Myc-Set1 (Dehe *et al*, 2006) and Prp22AID-FLAG (Mendoza-Ochoa *et al*, 2019). 250 ml of culture at an OD600 of 0.8 was harvested by centrifugation at 1000 x *g* and washed twice in ice-cold 1 X PBS. The cell pellet was re-suspended in 900 µl lysis buffer (50 mM Tris-HCl pH 7.5, 2 mM Mg2Cl2, 150 mM NaCl, 0.2% NP-40 and one complete EDTA-free proteinase inhibitor tablet (Roche #11836145001) and 400 µl zirconia beads. Cells were lysed using a Mini-Beadbeater-24 (BioSpec Products) twice at 2000 rpm for 2 min followed by 2 min on ice. The sample was centrifuged at 1000 x *g* for 2 min, the supernatant was collected and additionally centrifuged at 20,000 x *g* for 30 min at 4℃ and used for immunoprecipitation. The concentration of protein was measured using the Bradford assay, and 1 mg of protein used per IP. Prior to immunoprecipitation, extract was pre-cleared by adding ½ volume unconjugated Protein A/G Dynabeads (Life Technologies ##10001D/10003D). 50 µl Protein A/G Dynabeads conjugated to antibody were incubated with the appropriate volume of extract on a rotating wheel overnight at 4℃. The next day, beads were washed 8 times in lysis buffer (non-bound fraction kept for analysis). 20 µl of loading buffer was added to the beads, input and non-bound samples, which were boiled for 10 min before loading on a NuPAGE 4-12% Bis-Tris gel Bis-Tris (Invitrogen) and western blotting was performed.

### Set1 SUMOylation analysis

Cells were transformed with a plasmid encoding 6His-SUMO under the *CUP1* promoter (YEp352-6His-SUMO) or the corresponding empty vector (Niño *et al*, 2016). The transformed cells were grown on selective medium and stimulated overnight with 0.1 mM CuSO_4_. 200 OD_600_ of cells were collected and lysed with glass beads in a 20% TCA solution, the final TCA concentration was adjusted to 12%. Cell lysates were incubated at 4°C during 45 min, and precipitated proteins were collected by centrifugation. Proteins were resuspended in loading buffer (6 M guanidinium-HCl, 100 mM KH_2_PO_4_, 20 mM Tris-HCl, pH 8,0, 100 mM NaCl, 0,1% Triton X-100, and 10 mM imidazole). 6His-SUMOylated proteins were purified using Ni-NTA agarose beads (Qiagen), pre-equilibrated with loading buffer, and incubated for 1 h at room temperature. After incubation the beads were collected by centrifugation (3000 rpm, 2min) and washed twice with wash buffer (8 M urea, 100 mM Na_2_HPO_4_/NaH_2_PO_4_ pH 6,4, 10 mM Tris–HCl, pH 6.4, 10 mM Imidazole, 10 mM β-mercaptoethanol, and 0.1% Triton X-100). 6His-SUMOylated proteins were eluted with 50 ml 2X laemmli buffer (95°C, 5min). The proteins in the eluted fraction were analyzed by Western blot using anti-MYC (for endogenous Myc-Set1), anti-GAL4 (for GBD-Set1-full length and fragments) and polyclonal rabbit anti-Smt3.

### Interaction Snf2-AT-hook with SET1C and SET1C-762

GST-tagged Snf2 fragments (final concentration 250 nM) were combined with either the full-length SET1C or C762SET1C (final concentration 25 nM), along with 15 µl of glutathione-Sepharose 4B resin and BSA (final concentration 0.2 mg/ml) in a binding buffer containing 20 mM Tris-Cl (pH 7.9), 150 mM NaCl, 0.2 mM EDTA, 20% glycerol, 0.1% NP-40, and 1 mM PMSF, making a total volume of 300 µl. The mixtures were rotated at 4°C for 3 hours, followed by four washes with the binding buffer. The proteins bound to the resin were then eluted, separated by SDS-PAGE, and analyzed by immunoblotting.

### CO-IP of Snf2-Myc and Snf2ΔRG-Myc with Set1

WT cells or expressing *Snf2-Myc* and *Snf2ΔRG-Myc* (200 ml) were grown at 30°C in YPD to and O.D_600_ = 0.8. Cells were collected by centrifugation, washed once with 10 mM Tris-HCL, pH 8.0 and snap-frozen in liquid nitrogen. Cells were lysed using Retsch with the following parameters: 2 times 2 min at 30 m/S. The powder was resuspended in 3 ml of TMG 50 (10 mM Tris-HCL, pH 8.0, 0,1 mM MgCl_2_, 10% (V/V) glycerol, 50 mM NaCl, 0.1 mM EDTA, 0.1 mM DTT) containing protease inhibitors, 10 mM MG-132 and 1 mM PMSF. Cell lysates were clarified by centrifugation (13 300 rpm, 15’, 4°C) and 1.4 ml of supernatant were recovered. Protein concentration was determined using nanodrop and samples were adjusted to the same concentration by adding lysis buffer. Immunoprecipitation was performed overnight with 5 ml of 9E10 antibody (Santa Cruz Biotechnology) following by an incubation for 3 hours with 25 ml of pre-equilibrated protein-G dynabeads (Invitrogen). Immunoprecipitations were washed 3X with TMG 50 buffer and eluted using 25 ml of 1X Laemmli loading Buffer. Samples were resolved on a 7.5% Acrylamide gel, transferred on a nitrocellulose membrane and reveal with either anti-Myc 9E10 (Santa Cruz Biotechnology) and anti-Set1 antibodies.

### Snf2-GFP Complex purification

800 ml of *Snf2-GFP* and *Snf2-GFP set1Δ* cells were grown at 30°C in YPD to O.D_600_ = 0.8. Cell pellets were treated as described above except that supernatants were incubated with 25 ml of GFP nanobody coated Dynabeads (Gift from M. Modesti, CRCM Marseille) for 250 min, wash 3X with TMG 50 buffer and eluted in 30 ml of 2X Laemmli Loading buffer containing 50 mM DTT. For Western blotting, 1 ml was resolved on a NuPAGE 4-12% Bis-Tris Gel (Invitrogen), transferred on a nitrocellulose membrane and revealed using an anti-GFP antibody (A-11122-Invitrogen). For the Silver Staining, 2,5 ml were resolved on a NuPAGE 4-12% Bis-Tris Gel (Invitrogen) and stained using Pierce^TM^ Silver Stain kit (Thermo Scientific).

### Mass spectrometry-based identification of post-translational modifications in Snf2-GFP

Purified Snf2-GFP (n=3, biological replicates) from WT and *set1Δ* strains were loaded on NuPAGE™ 4–12% Bis–tris acrylamide gels according to the manufacturer’s instructions (Life Technologies). Running of protein was stopped as soon as proteins stacked in a single band. Protein containing bands were stained with Imperial Blue (Pierce), cut from the gel and digested with high sequencing grade trypsin (Promega) before mass spectrometry analysis. Briefly, gel pieces were washed and destained using few steps of 100 mM NH_4_HCO_3_. Destained gel pieces were shrunk with 100 mM ammonium bicarbonate in 50% acetonitrile and dried at RT. Protein spots were then rehydrated using 10mM DTT in 25 mM ammonium bicarbonate pH 8.0 for 45 min at 56°C. This solution was replaced by 55 mM iodoacetamide in 25 mM ammonium bicarbonate pH 8.0 and the gel pieces were incubated for 30 min at room temperature in the dark. They were then washed twice in 25 mM ammonium bicarbonate and finally shrunk by incubation for 5 min with 25 mM ammonium bicarbonate in 50% acetonitrile. The resulting alkylated gel pieces were dried at room temperature. The dried gel pieces were reswollen by incubation in 25 mM ammonium bicarbonate pH 8.0 supplemented with 12.5 ng/µl trypsin (Promega) for 1h at 4°C and then incubated overnight at 37°C. Peptides were harvested by collecting the initial digestion solution and carrying out two extractions; first in 5% formic acid and then in 5% formic acid in 60% acetonitrile. Pooled extracts were dried down in a centrifugal vacuum system. Samples were reconstituted in 0.1% TFA 4% acetonitrile before mass spectrometry using an Orbitrap Fusion Lumos Tribrid Mass Spectrometer (ThermoFisher Scientific, San Jose, CA) online with an Ultimate 3000RSLCnano chromatography system (ThermoFisher Scientific, Sunnyvale, CA). Peptides were separated at 40°C using a two steps linear gradient (4–20% acetonitrile/H2O; 0.1% formic acid for 110 min and 20-32% acetonitrile/H2O; 0.1% formic acid for 10 min). An EASY-Spray nanosource was used for peptide ionization (2,200 V, 275°C). MS was conducted using a data-dependent acquisition mode (DDA). The Orbitrap Lumos was used in data dependent mode to switch consistently between MS and MS/MS. Time between Masters Scans was set to 3 seconds. MS spectra were acquired with the Orbitrap in the range of m/z 400-1600 at a FWHM resolution of 120 000 measured at 400 m/z. AGC target was set at 4.0e5 with a 50 ms Maximum Injection Time. For internal mass calibration the 445.120025 ion was used as lock mass. The more abundant precursor ions were selected and collision-induced dissociation fragmentation was performed in the ion trap to have maximum sensitivity and yield a maximum amount of MS/MS data. Number of precursor ions was automatically defined along run in 3s windows using the “Inject Ions for All Available parallelizable time option” with a maximum injection time of 300 ms. The signal threshold for an MS/MS event was set to 5000 counts. Charge state screening was enabled to exclude precursors with 0 and 1 charge states. Dynamic exclusion was enabled with a repeat count of 1 and duration of 60 s.

For identification of PTMs the region corresponding to Snf2-GFP was excised from the gel and subjected to enzymatic digestion under the same conditions as above. Raw mass spectrometry files were analysed using MaxQuant software (version 1.6.3.4) as above with the exception of the Max Missed cleavage set at 5, the FDR at peptide and proteins levels at 5% and with the following variable modifications activated; Lysine acetylation (+42.0106); Serine, threonine and tyrosine phosphorylation (+79.966); Lysine and arginine methylation (+14.0157), dimethylation (+28.0313) and trimethylation (+42.0470).

### Data Processing Protocol

Relative intensity-based label-free quantification (LFQ) was processed using the MaxLFQ algorithm from the freely available MaxQuant computational proteomics platform, version 1.6.2.1 (Cox et al. 2014, Cox et al. 2008). The acquired raw LC Orbitrap MS data were first processed using the integrated Andromeda search engine (Cox et 2011). Spectra were searched against the Saccharomyces cerevisiae (organism ID 4932) extracted from UniProt containing 7904 entries. The following parameters were used for searches: (i) trypsin allowing cleavage before proline; (ii) two missed cleavage was allowed; (iii) cysteine carbamidomethylation (+57.02146) as a fixed modification and methionine oxidation (+15.99491) and N-terminal acetylation (+42.0106) as variable modifications. The match between runs option was enabled. The false discovery rate (FDR) at the peptide and protein levels were set to 1% and determined by searching a reverse database. For protein grouping, all proteins that cannot be distinguished based on their identified peptides were assembled into a single entry according to the MaxQuant rules. The statistical analysis was done with Perseus program (version 1.6.1.3) (Tyanova and Cox 2018) from the MaxQuant environment (www.maxquant.org). Quantifiable proteins were defined as those detected in above 70% of samples in one condition or more. Protein LFQ normalized intensities were base 2 logarithmized to obtain a normal distribution. Missing values were replaced using data imputation by randomly selecting from a normal distribution centered on the lower edge of the intensity values that simulates signals of low abundant proteins using default parameters (a downshift of 1.8 standard deviation and a width of 0.3 of the original distribution). The protein composition of the complex was determined using, a two-sample t-test using permutation-based FDR-controlled at 0.01 and employing 250 permutations and a scaling factor s0 with a value of 3. All proteins passing these criteria with a positive fold change are considered to be the potential interactome of Snf2.

### Antibodies

For IP: Mouse anti-FLAG (Sigma M2 #F1804); 10 μl/IP with Dynabeads and Protein G.

For westerns - primary antibodies: Mouse anti-c-MYC (Santa Cruz #SC-40x) 1:1000; Rat anti-FLAG (Agilent #200474) 1:1000; Mouse anti-Gal4 Binding Domain (Euromedex) 1:5000; Rabbit anti-Smt3 is a gift from B. Palancade (Institut Jacques Monod, Paris). Anti-Bre2 is a gift from Peter Nagy. The rabbit anti-Spp1 antibody was developed in V. Géli’s lab. The anti-Spp1 antibody is showing significant cross-reactivity with other cellular proteins but did allow the identification of Spp1 in purified and partially purified fractions. Secondary antibodies: Goat anti-mouse IRDye680RD (LI-COR #926-68070) 1:10,000, Goat anti-rat IRDye800RD (LI-COR #926-32219) 1:10,000.

## DATA AVAILABILITY

Additional data and reagents are available upon request to the corresponding authors. The mass spectrometry proteomics data have been deposited to the ProteomeXchange Consortium (www.proteomexchange.org) via the PRIDE partner repository (https://www.ebi.ac.uk/pride/login): Snf2-B3 data (accession number PXD061448), Snf2-GFP complex (accession number PXD061496), Snf2 PTM (accession number PXD061531).

## ACKNOWLEDGEMENTS

We are grateful to Sandra Holbein and Andre Halbach for help with MBP-Nrm1 fusion proteins. We thank Benoit Palancade for his help in the interpretation of the Y2H results and reagents.

## AUTHOR CONTRIBUTIONS

B.D. J. K and V.G. designed the study and secured the funding. All the Y2H screens and prey identification were performed by Hybrigenics in close interaction with B.D. and V. G. P.L. performed molecular cloning, contributed to validation experiments, and purified Snf2-GFP for subsequent analyses. SET1C and SET1C-C762 purification and interaction assays with Snf2-AT-hook were performed by J. K. and K.P. Set1 SUMOylation experiments were performed by C. N. M. D and B. D. organized the results shown in Table S2. L. L constructed the F1-F5 fragments and contributed to validation experiments. Pull-down experiments and Nrm1 methylation assay were performed by B. D. TAP-Nis1 data were provided by M.O. I.M. and J. B. performed and discuss the Set1/Prp22 interaction. D.K. P and H. H. K performed mass spectrometry analysis of the Snf2-B3. S.A. and L.C. performed mass spectrometry analysis of the Snf2-GFP. V.G. and B. D. wrote the manuscript with inputs of the authors.

