## Supplementary information for "A systematic interactome of SET1C expands its functional landscape and identifies candidate regulatory connections"

#### Tables

**Table S1. Plasmids and yeast strains used in this study.**

**Table S2. All SET1C interactors identified in the 10 Y2H screens.**
The interactors common to several subunit and to common Set1 fragments are shown on sheet 13 of the table.

**Table S3. Mass spectrometry analysis of TAP-Nis1 affinity purification**

**Table S4. H3K4-like domain proteins.** We used the scansite search algorithm (http://scansite.mit.edu) and systematically identified sequence motifs that are related to the Set1C modification site in histone H3.

**Table S5.** Snf2-GFP complex in WT and *set1Δ* cells

**Table S6.** Snf2 post-translational modifications in either WT or *set1*Δ strains

### Table S1

| PLASMIDS | DESCRIPTION | Source |
|---|---|---|
| pP6 | ADH1 promoter driving GAL4 activation domain, LEU2, 2μ ori, AmpR | Hybrigenics |
| pB66 | ADH1 promoter driving GAL4 DNA binding domain, TRP1, 2μ ori, KanR | Hybrigenics |
| pB43 | ADH1 promoter driving LEXA DNA binding domain, TRP1, 2μ ori | Hybrigenics |
| pB66-FL SET1 | SET1 into pB66 | Hybrigenics |
| pB66-FL SET1 1-754 | SET1 1-754 into pB66 | Hybrigenics |
| pB66- SET1 754-1081 | SET1 754-1081 into pB66 | Hybrigenics |
| pB66-BRE2 | BRE2 into pB66 | Hybrigenics |
| pB27-SWD1 | SWD1 into pB27 | Hybrigenics |
| pB27-SWD3 | SWD3 into pB27 | Hybrigenics |
| pB27-SDC1 | SDC1 into pB27 | Hybrigenics |
| pB27-SGH1 | SHG1 into pB27 | Hybrigenics |
| pB27-SPP1 | SPP1 into pB27 | Hybrigenics |
| pB43-SWD2 | SWD2 into pB43 | Hybrigenics |
| pB66-Set1 F1 | ADH1 promoter driving GAL4 DNA binding domain-SET1 F1 (aa 1-236), TRP1, 2μ ori, KanR | Lara Lee (CRCM) |
| pB66-Set1 F2 | ADH1 promoter driving GAL4 DNA binding domain-SET1 F2 (aa 240-586), TRP1, 2μ ori, KanR | Lara Lee (CRCM) |
| pB66-Set1 F3 | ADH1 promoter driving GAL4 DNA binding domain-SET1 F3 (aa 590-769), TRP1, 2μ ori, KanR | Lara Lee (CRCM) |
| pB66-Set1 F4 | ADH1 promoter driving GAL4 DNA binding domain-SET1 F4 (aa 770-945), TRP1, 2μ ori, KanR | Lara Lee (CRCM) |
| pB66-Set1 F5 | ADH1 promoter driving GAL4 DNA binding domain-SET1 F5 (aa 946-1080), TRP1, 2μ ori, KanR | Lara lee (CRCM) |
| pB66-Set1 F3+F4 | ADH1 promoter driving GAL4 DNA binding domain-SET1 F5 (aa 590-945), TRP1, 2μ ori, KanR | Lara Lee (CRCM) |
| YEp352-6His-SUMO | 6His-SUMO under the CUP1 promoter, URA3 | Benoit Palancade (IJM) |

| STRAINS | GENOTYPE | Source |
|---|---|---|
| Y187 | MATa Gal4D Gal80D ade2-101 his3 leu2-3,-112 trp1-901 ura3-52 URA3::UASGAL1-LacZ, (met-) | |
| L40ΔGAL4 | MATa, his3-A200, trp1-901, leu2-3,112, ade2, LYS2::(1exAop)4—HIS3, URA3::(1exAop)8-LacZ, ga180) ΔGAL4 | |
| YHGX13 | Mat alpha, idem Y187 + ade2-101::loxP-kanMX-loxP | |
| HF7c | MATa Gal4-452 Gal80-538 ade2-101 his3-D200 leu2-3,112 trp1-901 ura3-52 lys2-801 URA3::Gal4 17mers (X3)-CyC1TATA-LacZ lys2::GAL1UAS-GAL1TATA-HIS3 | |
| CG1945 | Mata idem HF7c + cyhR | |
| TOTO | Diploid strain resulting from mating between CG1945 and YHGX13 | |
| W303 | MATa ade2-1 leu2-3,112 trp1-1 can1-100 ura3-1 ade2-1 his3-11,15 | |
| W303-Myc-Set1::TRP | MATa leu2-3,112 trp1-1 can1-100 ura3-1 ade2-1 his3-11,15 MYC-SET1::TRP | Dehe *et al.* 2006 |
| W303-Prp22-AID*-6FLAG_pFUI_PADH1-409-TIR1 | MATa leu2-3,112 trp1-1 can1-100 ura3-1 ade2-1 his3-11,15PADH1-409-Os TIR1-NatMX PRP22::PRP22-AID*-6FLAG-HygMX | Mendoza-Ochoa *et al*. 2018 |
| W303-Prp22-AID*-6FLAG_pFUI_PADH1-409-TIR1 Myc-Set1::TRP | MATa leu2-3,112 trp1-1 can1-100 ura3-1 ade2-1 his3-11,15PADH1-409-Os TIR1-NatMX PRP22::PRP22-AID*-6FLAG-HygMX [pRS426-FUI1-URA3 MYC-SET1::TRP | This study |
| W303-Snf2-Myc::KAN | MATa ade2-1 leu2-3,112 trp1-1 can1-100 ura3-1 ade2-1 his3-11,15 SNF2-MYC::KanMX | This study |
| W303- *snf2ΔRG-Myc::KAN* | MATa ade2-1 leu2-3,112 trp1-1 can1-100 ura3-1 ade2-1 his3-11,15 *snf2ΔRG-MYC::KanMX* | This study |
| BY4741 | MATa his3Δ1 leu2Δ0 met15Δ0 ura3Δ0 | B.Palancade |
| BY4741-Snf2-GFP::HIS3 | MATa his3Δ1 leu2Δ0 met15Δ0 ura3Δ0 SNF2-GFP::HIS3 | B. Palancade |
| BY4741-Snf2-GFP::HIS3 *SET1::NAT* | MATa his3Δ1 leu2Δ0 met15Δ0 ura3Δ0 SNF2-GFP::HIS3 *SET1::NatMX* | This study |

### SUPPLEMENTARY FIGURES

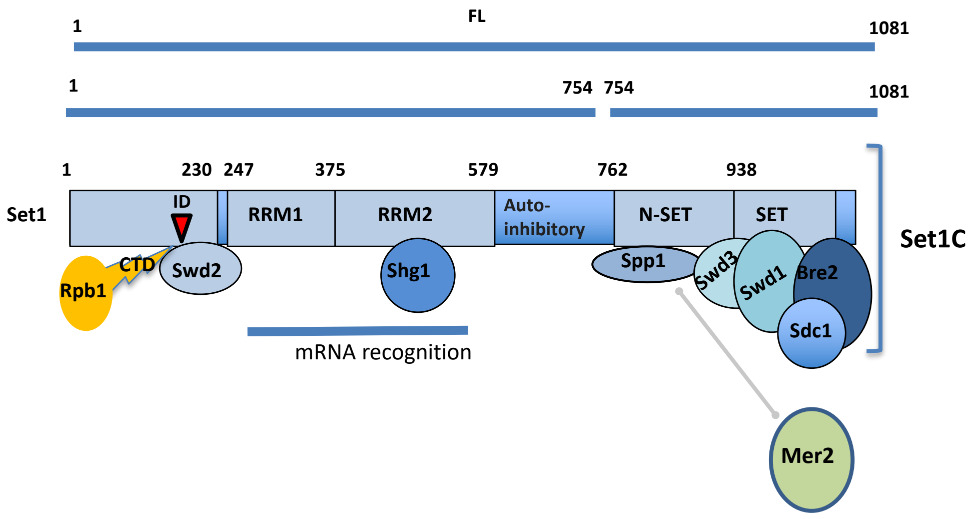


**Figure S1. Schematic representation of SET1C/COMPASS.** These Y2H screens revealed the interaction of Mer2 with Spp1(Acquaviva *et al.* 2013) and of Rbp1-CTD with the N-terminal region of Set1 (Bae *et al.* 2020).

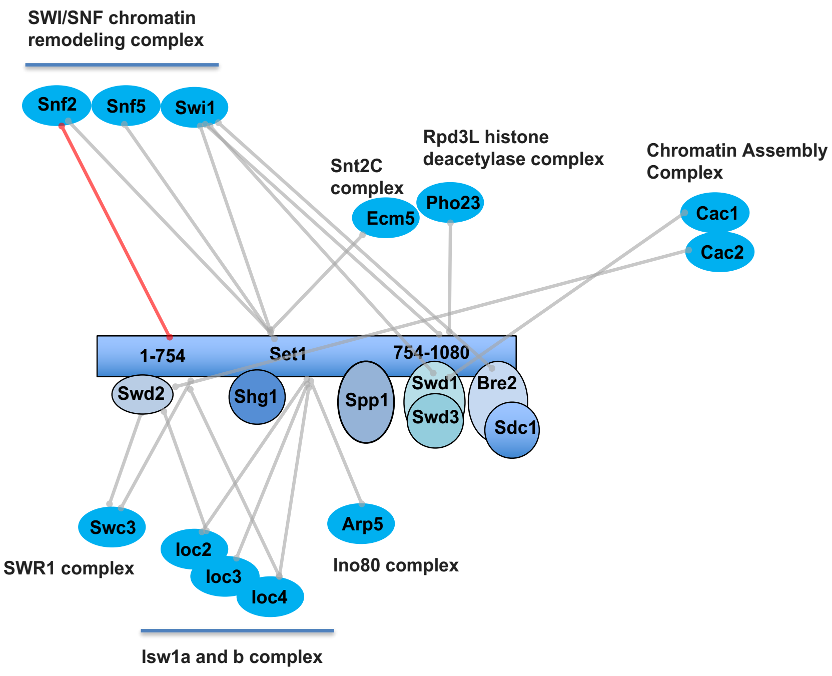


**Figure S2. Chromatin regulators identified in all the two-hybrid screens.** The lines indicate the individual proteins involved in the Y2H interaction. Red line refers to a very high confidence Y2H interaction. All Y2H interactors are described in Table S2. Interactors are grouped according to the complex to which they belong.

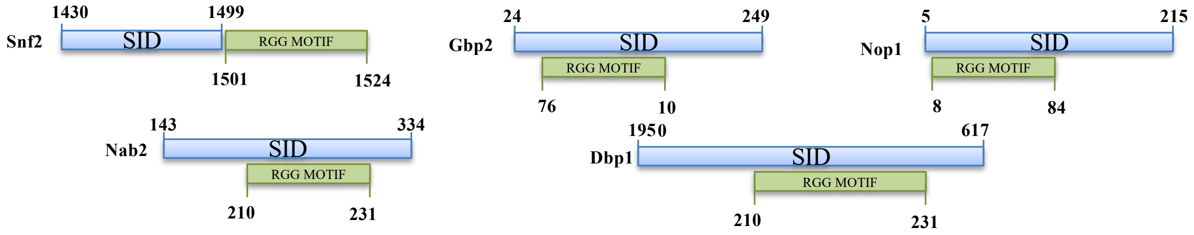


**Figure S3.** SID (blue) and RGG motif (green) within the RGG proteins. The interaction domains indicated represent the minimal overlapping DNA sequence present in multiple independent Y2H interacting clones of the same gene. Each genomic fragment of a Y2H clone was analyzed, and the shared overlapping region for given gene was determined to be the only common element among all interacting clones. As such, this region represents the minimal sequence required for interaction.

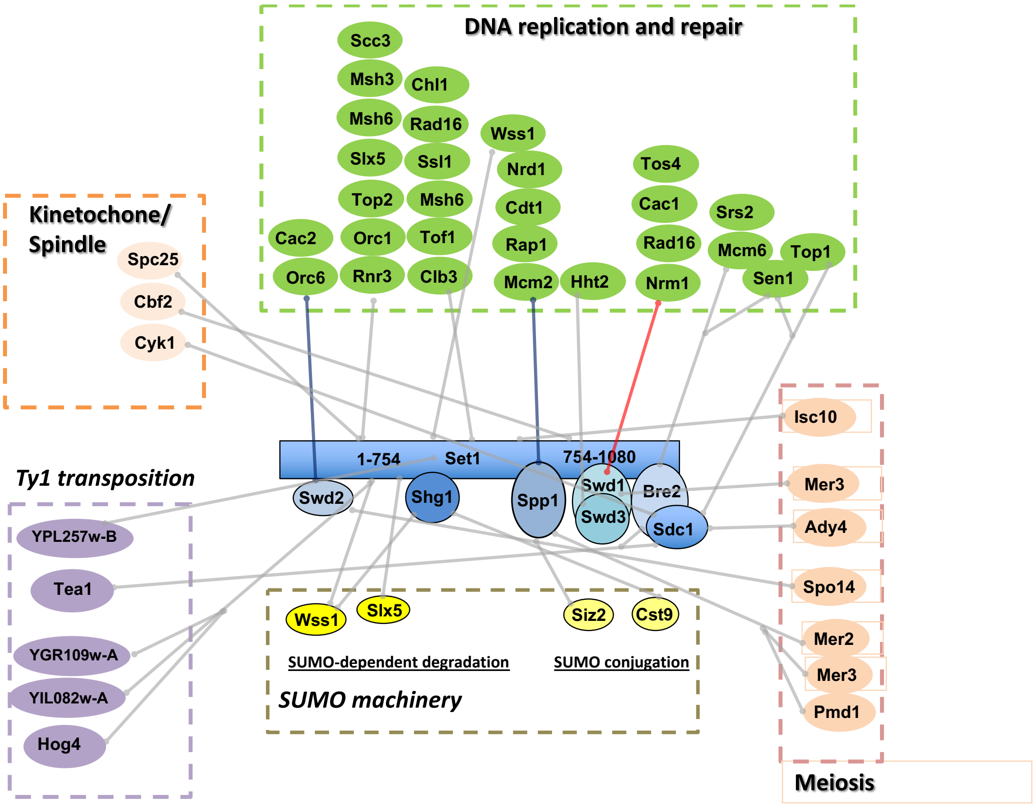


**Figure S4.** SET1C Y2H interactors involved in DNA transactions.
The processes in which Y2H interactors are involved are shown in the figure. Red and blue line refers to the confidence of the Y2H interaction.

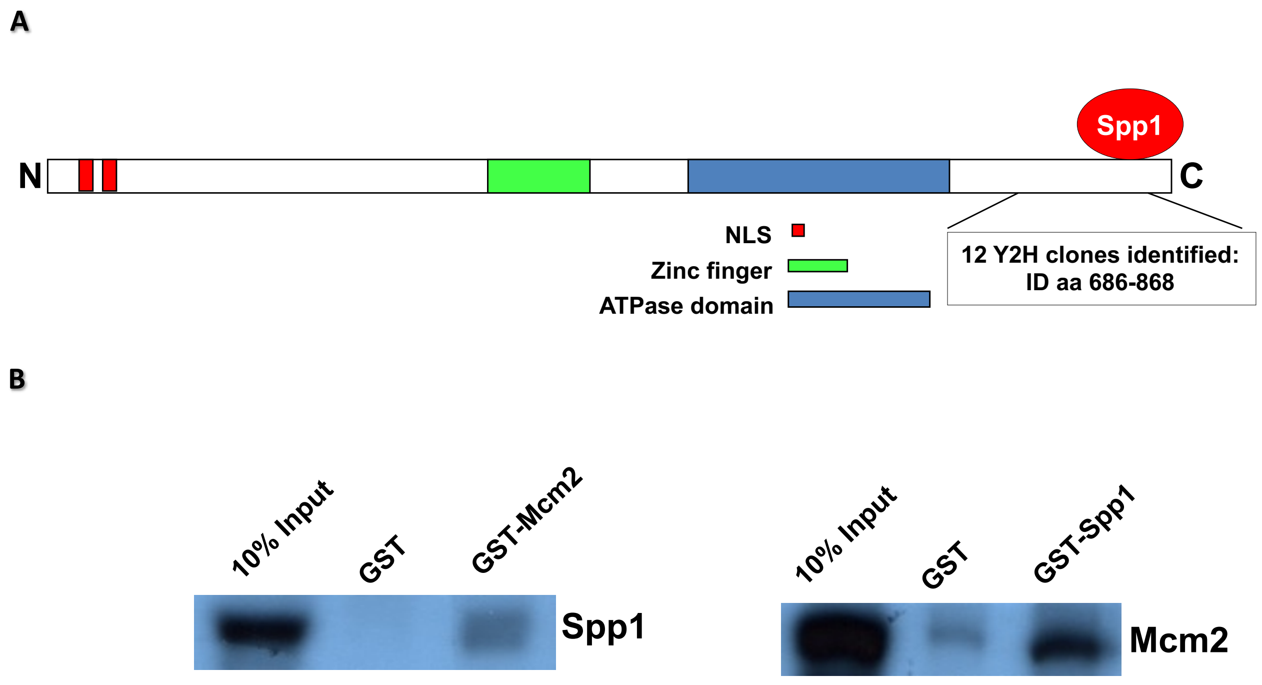


**Figure S5. Spp1 interacts weekly with Mcm2 *in vitro*.**
**A**) Schematic representation of Mcm2. The minimal region of interaction with Spp1 is located at the extreme C-terminus of Mcm2. **B**) Pull-down assays were performed with GST, GST-Spp1 and GST-Mcm2 fusion proteins. $^{35}$S-labelled *in vitro* translated proteins served as preys; input shows 10% of radioactive material included in binding reactions.

| **protein** | **size (kDa)** | **motif** | **function** |
|---|---|---|---|
| Histone H3 | 35 | **ARTKQT** | core histone protein required for chromatin assembly |
| Nrm1 | 57 | **KTKQT** | transcriptional co-repressor of MBF-regulated gene expression |
| Nrd1 | 84 | **RSKQ** | transcription termination and 3' end maturation of nonpolyadenylated RNAs |
| Rix1 | 107 | **KTKQS** | processing of ITS2 sequences from 35S pre-rRNA |
| Dbf2 | 86 | **RTKQ** | transcription and stress response, part of a network of genes in exit from mitosis |
| Dbf20 | 86 | **RTKQ** | late nuclear division, one of the mitotic exit network (MEN) proteins |
| Not5 | 86 | **KTKQ** | transcription initiation, elongation and in mRNA degradation |
| Mcm2 | 120 | **ARTK** | DNA replication |
| Prp8 | 300 | **RTKQ** | second catalytic step of splicing; mutations of human Prp8 cause *retinitis pigmentosa* |

**Figure S6.** H3K4-like proteins. Shown are selected proteins containing sequences similar to the modification site found in histone H3. An exhaustive list can be found in Table S4.

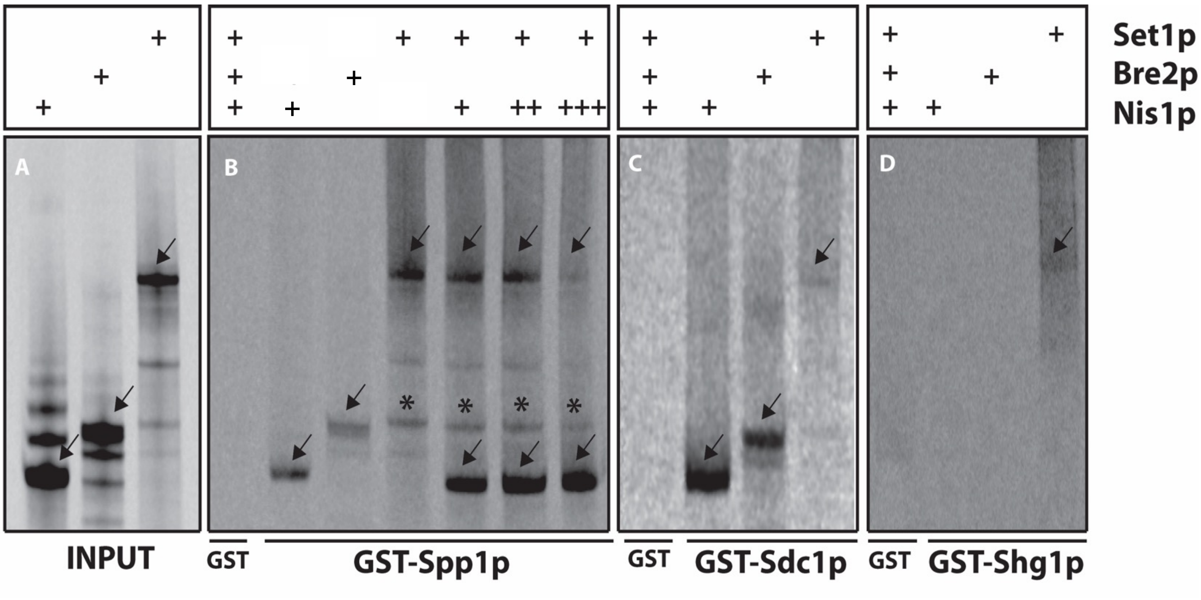


**Figure S7. Spp1 and Sdc1 interacts with Nis1 *in vitro***

Pull-down assays were performed with GST, GST-Spp1, GST-Sdc1 and GST-Shg1 fusion proteins as indicate at the bottom of the panels. $^{35}$S-labelled *in vitro* translated proteins served as preys; input shows 10% of radioactive material included in binding reactions. The asterisk is marking a band that is an undefined Set1 in vitro translation product and it is marked so that it is not confused with full-length Bre2. The arrows point to the expected full-length in vitro translation products. Nis1 (46 KDa), Bre2 (58 KDa), Set1 (124 KDa).

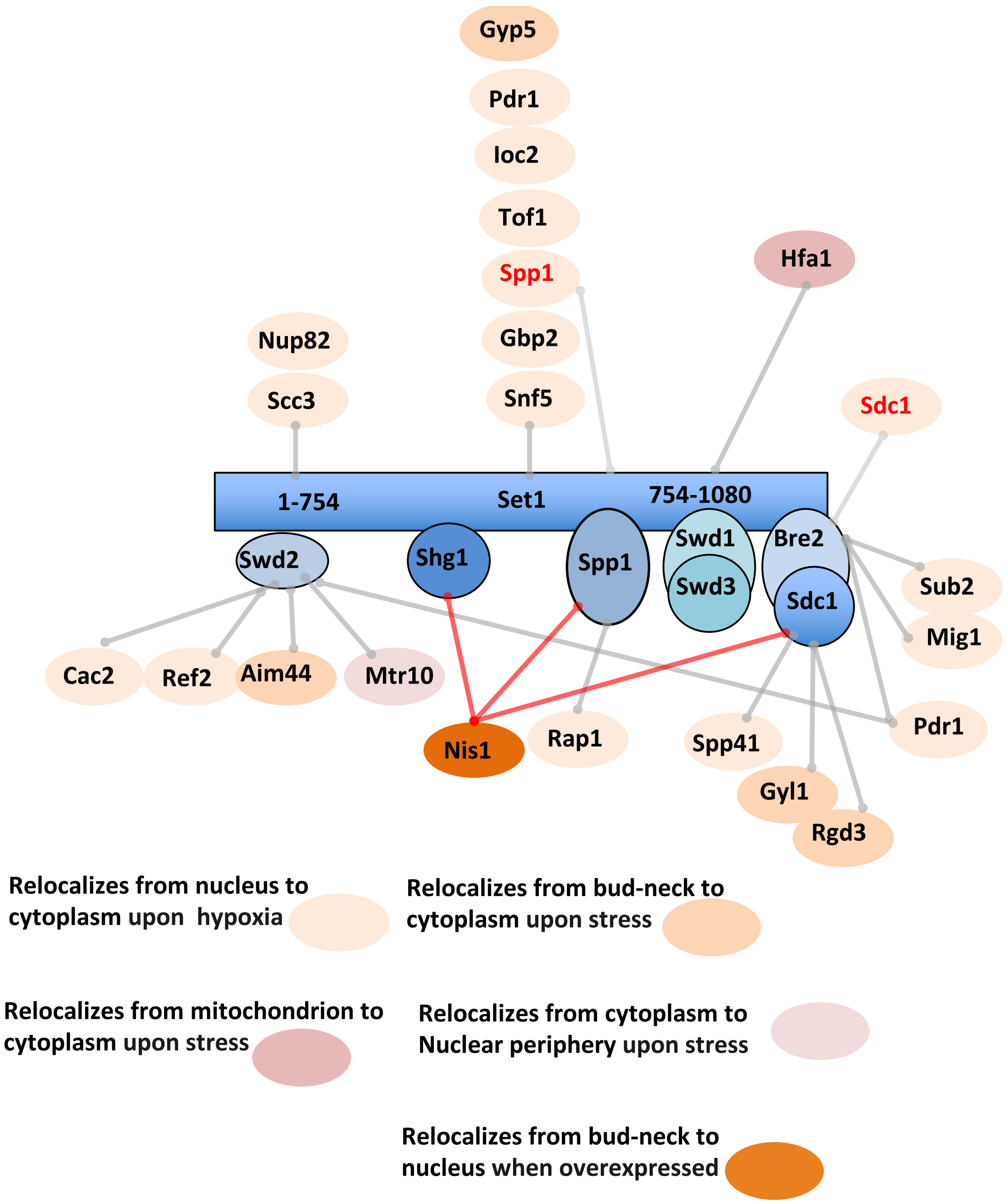


**Figure S8. Identification of SET1C Y2H interactors showing changes in localization following hypoxia.**
SET1C interactors that change localization are shown. In hypoxia, 243 proteins change their localization (Henke *et al.* 2011). Several components of the SWI/SNF complex were found to rapidly change location (from nucleus to cytosol) in hypoxia (Dastidar *et al.* 2012).

**A**

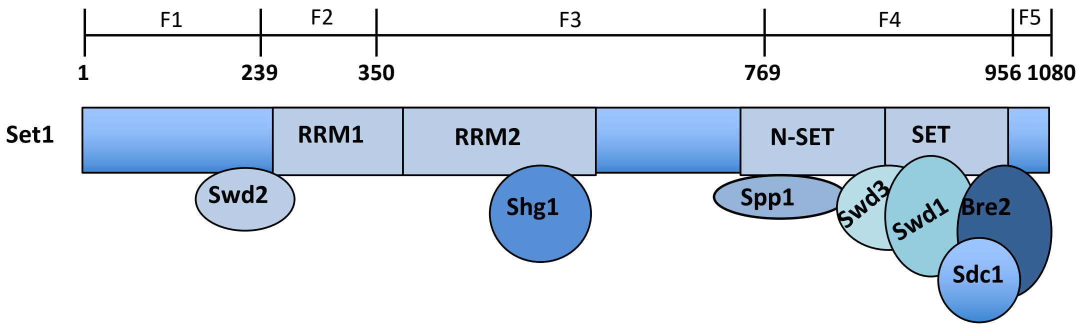


**B**

| POSITION | PEPTIDE | SCORE | CUTOFF | TYPE |
|---|---|---|---|---|
| 156 | FHYFDPI**K**GEFFNKD | 43.08 | 16 | SUMOylation |
| 332 | GCFIMGF**K**FEVILNK | 18.832 | 16 | SUMOylation |
| 356 | KFVEINV**K**KLQKLQE | 38.727 | 36.625 | SUMOylation |
| 521 | NSTNVPI**K**YESKEEF | 43.49 | 16 | SUMOylation |
| 698 | EAPDKKF**K**SESEPTT | 19.129 | 16 | SUMOylation |
| 769 | MDLQNAI**K**DEEDMLI | 46.442 | 16 | SUMOylation |
| 836 | LQPGSSF**K**AEGFRKI | 16.286 | 16 | SUMOylation |
| 1060 | DYKFERE***K*** DDEERLP | 42.152 | 36.625 | SUMOylation |

**Figure S9. Set1 SUMOylation analysis.**

**A**) Schematic representation of the F1-F5 Set1 fragments. **B**) List of predicted SUMOylated sites by GPS-SUMO 1.0 (Zhao *et al.* 2014). The predicted SUMOylated lysine is bold underlined.

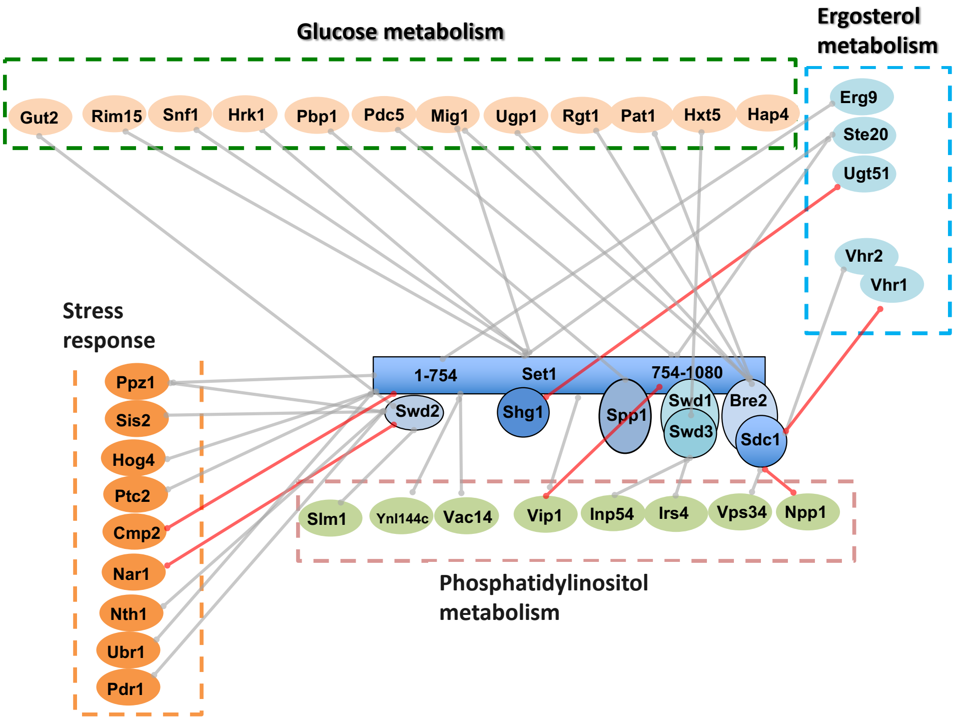


**Figure S10. SET1C Y2H interactors are involved in stress responses and metabolism pathways.** The processes in which Y2H interactors are involved are shown in the figure. Red arrows indicate high confidence Y2H interactions.

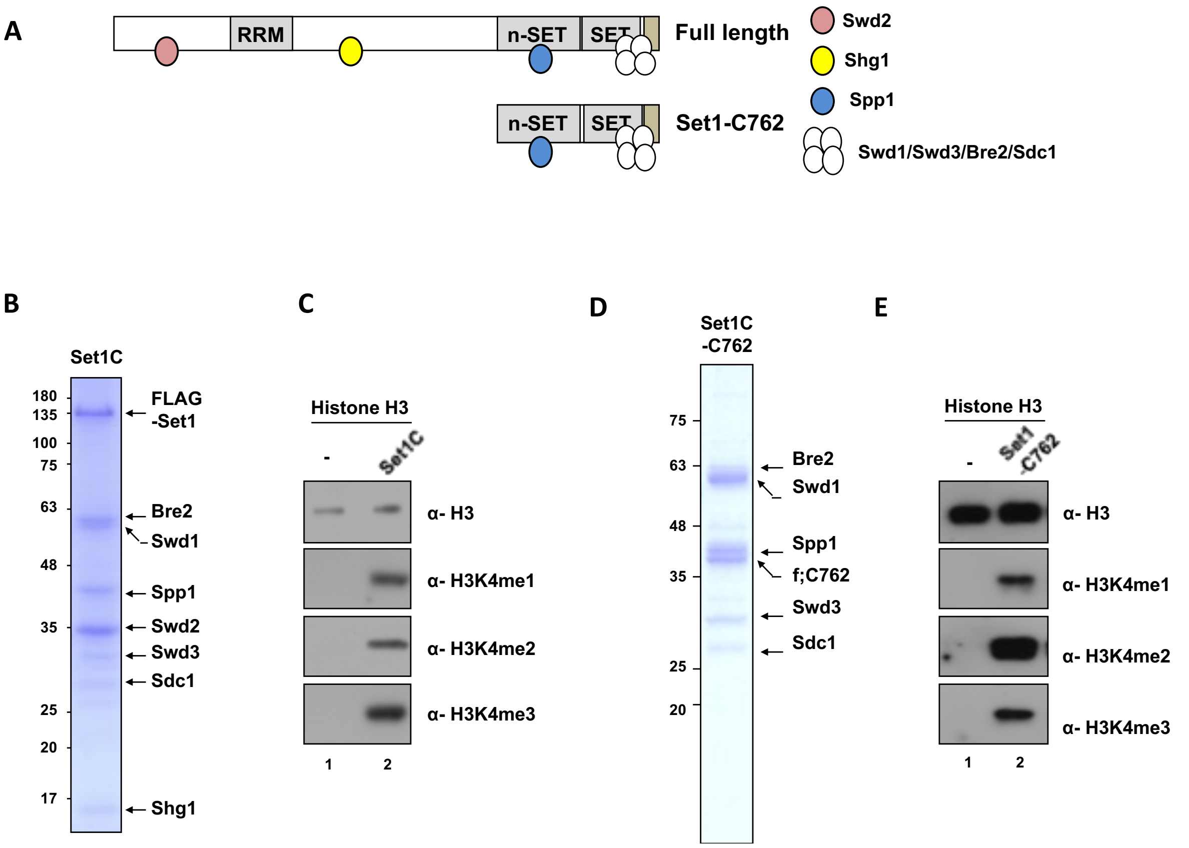


**Figure S11. Purification and activity of SET1C and SET1C-C762.**

**A**) A schematic diagram of the subunits used for the reconstitution of SET1C and Set1-C762 complexes. **B**) SDS-PAGE/Coomassie Blue staining of purified SET1C. **C**) *In vitro* methyltransferase assay using SET1C and free histone H3. **D**) SDS-PAGE/Coomassie Blue staining of purified Set1-C762 complex. **E**) *In vitro* methyltransferase assay using Set1-C762 complex and free histone H3.

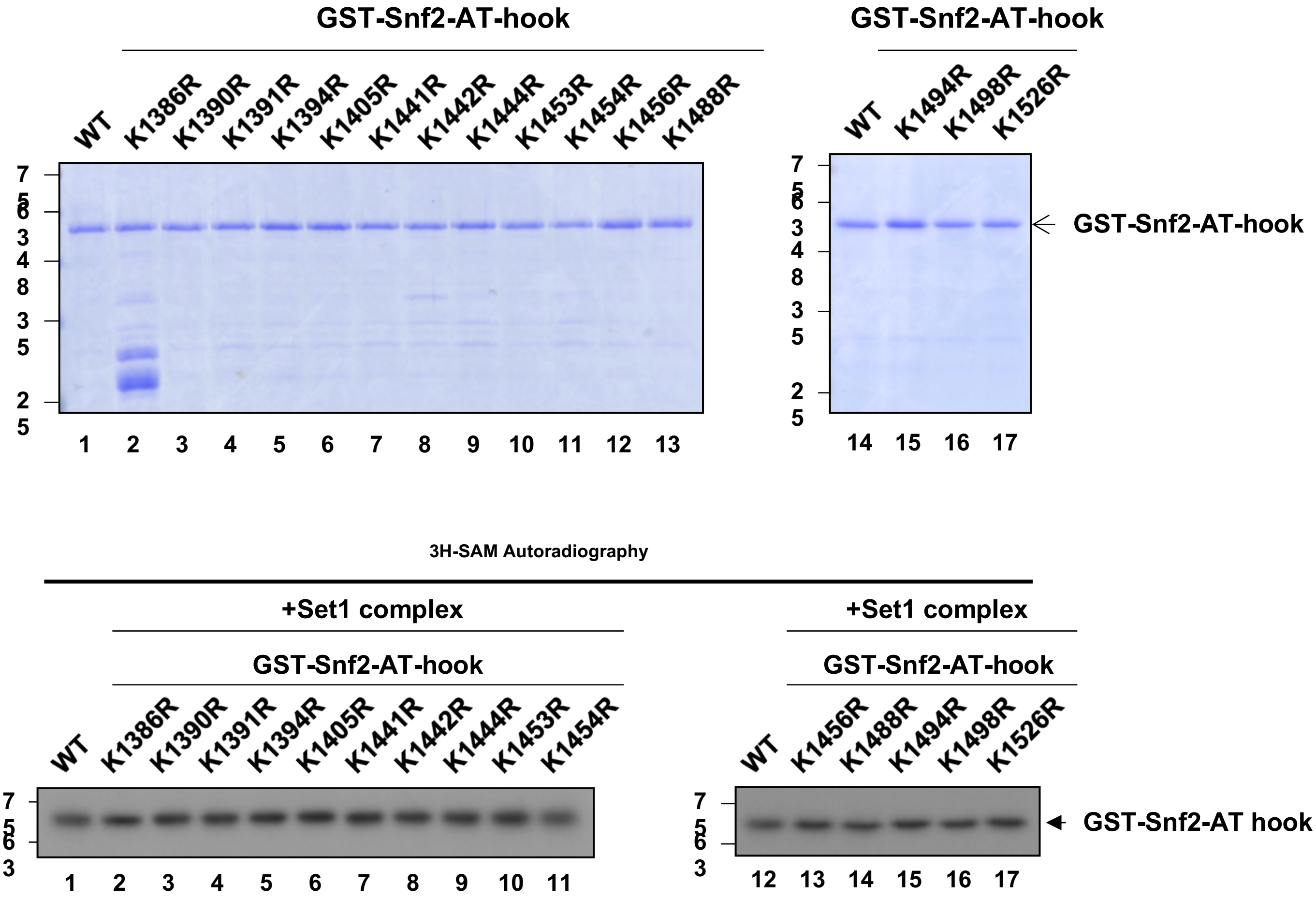


**Figure S12. 15 lysines of Snf2-AT-hook were replaced with arginine one by one.** Purified Snf2-AT-hook wild type and 15 mutant proteins were subject to Coomassie staining (upper) and *in vitro* methyltransferase assay with SET1C (lower).

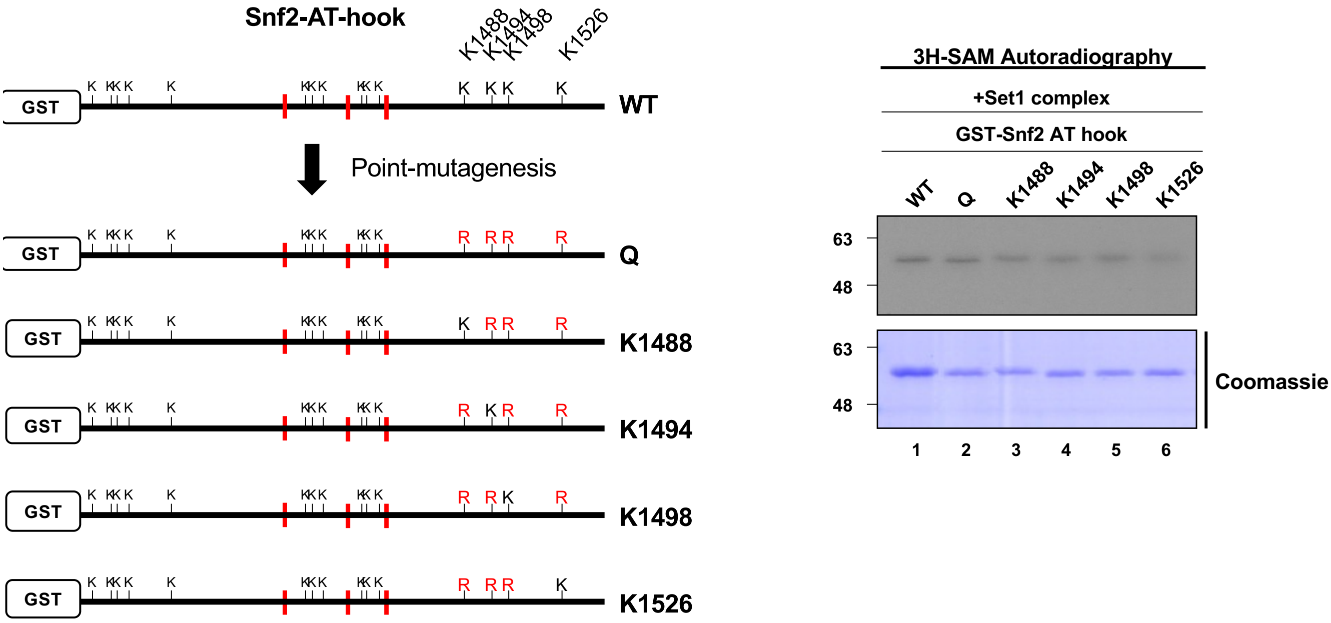


**Figure S13. A**) Designed AT-hook mutants with quadra mutation of all four lysines (K1488, K1494, K1498, K1526) or triple mutations among these four lysines. K1494 and K1498 are known to be acetylated by Gcn5. **B**) *In vitro* methyltransferase assay using reconstituted SET1C and Snf2-AT-hook mutants.

# A

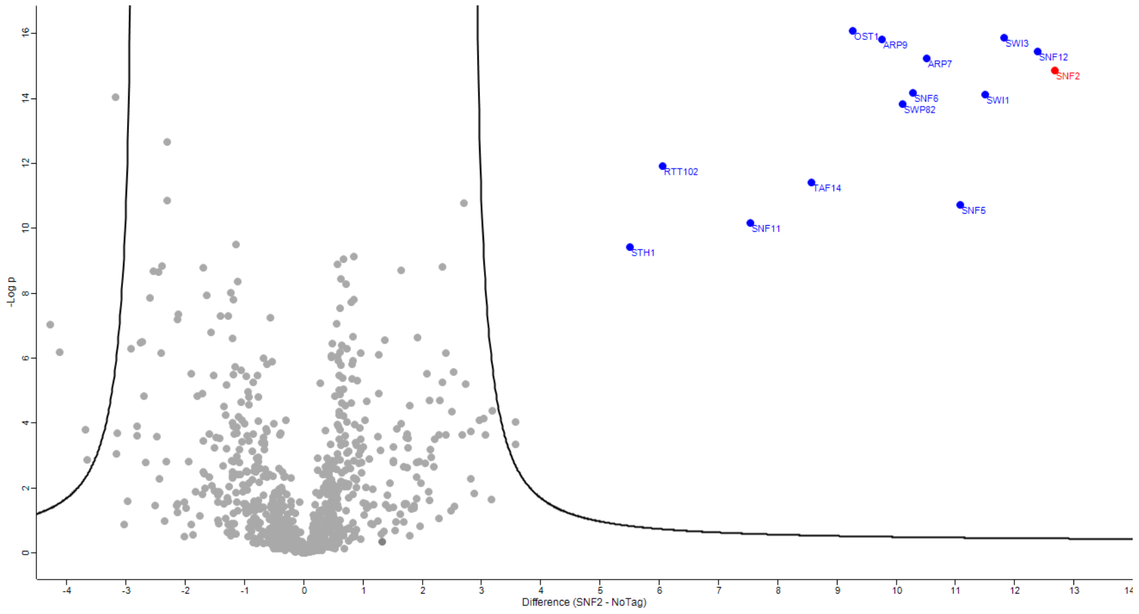


# B

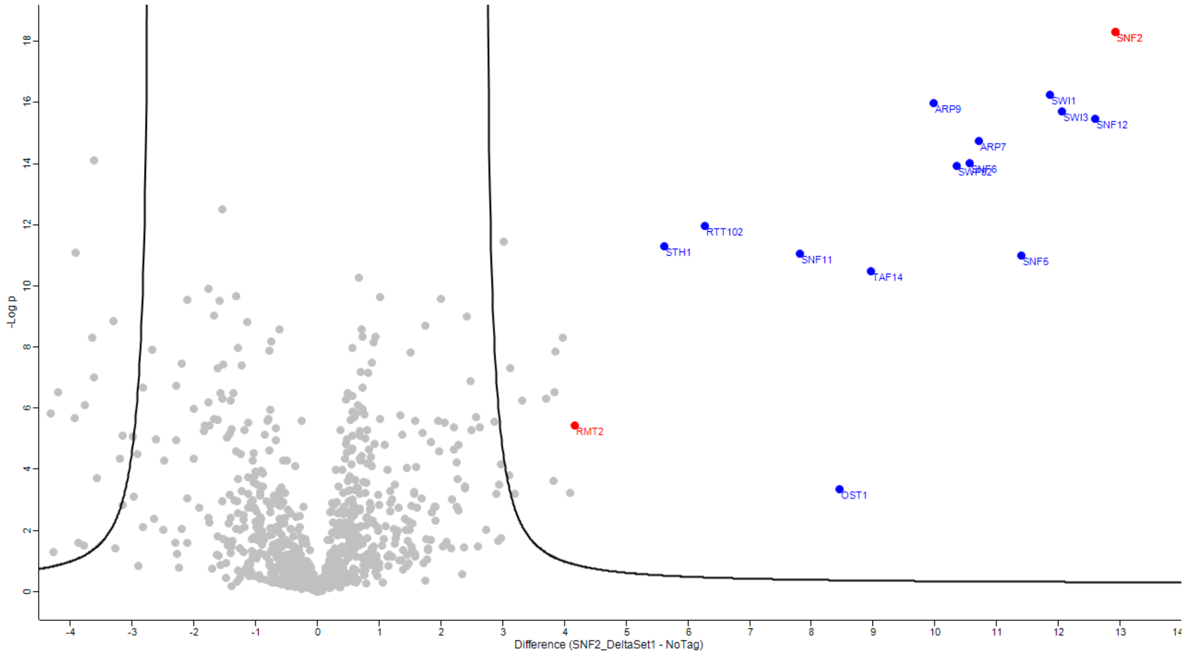


**Figure S14. Volcano plot describing the enrichment of proteins in the purifications of Snf2-GFP compared to the control (No tag) (see Table S5) created by Perseus.** A) In a wild-type strain, B) in the *set1Δ* strain. The curve represents the significant limit of quantification (S0 = 5 and FDR = 0.01).

WT

| Amino Acid | Peptide sequence and PTM localization probabilities | Position in peptide | Positions within the protein | PTM |
|---|---|---|---|---|
| K | LEGSENSEPPALESSPVTGDNSPSEDFMDIPK(0.5)PR | 32 | 1488 | Dimethyl |
| R | LEGSENSEPPALESSPVTGDNSPSEDFMDIPKPR(0.5) | 34 | 1490 | Dimethyl |
| K | TAGK(1)TSVKSAR | 4 | 1494 | Acetyl |
| K | TAGK(0.999)TSVKSARTSTRGR | 4 | 1494 | Methyl |
| S | TAGKT(0.723)SVKSARTSTR | 5 | 1495 | Phospho |
| S | TAGKTS(0.786)VKSARTSTR | 6 | 1496 | Phospho |
| K | TAGKTSVK(1)SAR | 8 | 1498 | Acetyl |
| K | TAGKTSVK(0.951)SARTSTR | 8 | 1498 | Dimethyl |
| K | TAGKTSVK(0.874)SARTSTR | 8 | 1498 | Methyl |
| S | TAGKTSVKS(0.9)ARTSTRGR | 9 | 1499 | Phospho |
| R | TAGKTSVKSAR(0.974)TSTRGR | 11 | 1501 | Methyl |
| S | TAGKTSVKSART(0.906)STRGR | 12 | 1502 | Phospho |
| S | TAGKTSVKSARTS(0.94)TR | 13 | 1503 | Phospho |
| T | TAGKTSVKSARTST(0.952)RGR | 14 | 1504 | Phospho |
| R | TSTR(0.999)GRGRGR | 4 | 1505 | Methyl |
| R | TAGKTSVKSARTSTR(0.477)GR | 15 | 1505 | Dimethyl |
| R | TSTRGR(0.944)GRGR | 6 | 1507 | Methyl |
| R | TAGKTSVKSARTSTRGR(0.477) | 17 | 1507 | Dimethyl |
| R | AR(0.496)NGLDYVRTPAAATSPIDIR | 2 | 1528 | Dimethyl |
| Y | ARNGLDY(0.966)VRTPAAATSPIDIR | 7 | 1533 | Phospho |
| R | ARNGLDYVR(0.504)TPAAATSPIDIR(1) | 9 | 1535 | Dimethyl |
| S | ARNGLDYVRTPAAATS(0.989)PIDIR | 16 | 1542 | Phospho |
| R | ARNGLDYVRTPAAATSPIDIR(1) | 21 | 1547 | Dimethyl |

*set1Δ*

| Amino acid | Peptide sequence and PTM localization probabilities | Position in peptide | Positions within proteins | PTM |
|---|---|---|---|---|
| R | PR(1)TAGKTSVK | 2 | 1490 | Methyl |
| T | T(0.609)AGKTSVKSARTSTR | 1 | 1491 | Phospho |
| K | TAGK(1)TSVKSARTSTR | 4 | 1494 | Acetyl |
| K | PRTAGK(0.835)TSVK | 6 | 1494 | Methyl |
| T | PRTAGKT(0.997)SVK | 7 | 1495 | Phospho |
| S | PRTAGKTS(0.998)VK | 8 | 1496 | Phospho |
| K | TAGKTSVK(1)SARTSTR | 8 | 1498 | Dimethyl |
| K | TAGKTSVK(1)SARTSTR | 8 | 1498 | Acetyl |
| S | TAGKTSVKSARTS(0.955)TR | 13 | 1503 | Phospho |
| T | TAGKTSVKSARTST(0.989)RGR | 14 | 1504 | Phospho |
| R | AR(1)NGLDYVRTPAAATSPIDIR | 2 | 1527 | Methyl |
| Y | ARNGLDY(0.803)VRTPAAATSPIDIR | 7 | 1532 | Phospho |
| R | ARNGLDYVR(1)TPAAATSPIDIR | 9 | 1534 | Dimethyl |
| T | TPAAAT(0.687)SPIDIR | 6 | 1540 | Phospho |
| S | ARNGLDYVRTPAAATS(0.984)PIDIR | 16 | 1541 | Phospho |
| R | ARNGLDYVRTPAAATSPIDIR(0.914) | 21 | 1546 | Methyl |
| K | EK(0.5)VAKQALDLYHFALNYENEAGR | 2 | 1549 | Dimethyl |
| K | EK(0.5)VAKQALDLYHFALNYENEAGR | 2 | 1549 | Trimethyl |

**Figure S15. Identified peptides showing localization probabilities from the amino sequence flanking the RG repeats (1485-1549).**

Peptides identified after Snf2-GFP digestion for wild type (WT) and *set1Δ* are indicated with their identified PTM. The red numbers indicate the probability of localization according to the MS2 peaks for the preceding amino acid.
